# Amyloid-β-regulated gene circuits for programmable Alzheimer’s disease therapy

**DOI:** 10.1101/2025.03.12.642808

**Authors:** Madeline R. Spetz, Hyosung Kim, Alexander P. Ligocki, Swathi R. Shekharan, Hannah J. Brien, Daniel Chavarria, Dylan J. Conger, Rebecca Shattuck-Brandt, Alena Shostak, Rebecca J. Embalabala, Matthew S. Schrag, Ethan S. Lippmann, Jonathan M. Brunger

**Author notes:** **Corresponding author** Correspondence to Jonathan M. Brunger.

## Abstract

Alzheimer’s disease (AD) is a neurodegenerative disease characterized in part by the accumulation of amyloid-β (Aβ). Monoclonal antibodies (mAbs) that target Aβ for clearance from the brain are FDA-approved; however, these therapies are associated with serious side effects, and their cognitive benefit remains debated. We present a potential engineered therapy for AD in which we enlist cells of the central nervous system as programmable agents for sculpting the neurodegenerative niche to mitigate glial reactivity and neuronal loss. We constructed a suite of Aβ-sensitive synthetic Notch (synNotch) receptors from anti-Aβ mAbs and show that cells expressing them can recognize distinct Aβ species with differential sensitivity. We validate Aβ-synNotch receptor activity in microglia and astrocytes, cells native to the brain that become dysfunctional in AD. SynNotch astrocytes engineered to express antagonists of the pro-inflammatory cytokines TNF and IL-1 resist a cytokine-induced, reactive phenotype. Finally, Aβ-synNotch cells potently upregulate transgene expression in response to Aβ deposits in 5xFAD mouse brains. We also adapt the Aβ receptor for a gene therapy, supporting a translational strategy that bypasses the need for *ex vivo* cell engineering. Together, these results establish Aβ-responsive synthetic receptors for conditional delivery of disease-modifying factors to the AD microenvironment.

## Introduction

Alzheimer’s Disease (AD) is a progressive, neurodegenerative disorder. The pathophysiology underlying AD is complex, though it is most often characterized by accumulation of amyloid-β (Aβ) and tau aggregates.^1–3^ These proteins have been the subject of intensive research in AD therapeutic development. Much of this work has focused on anti-Aβ immunotherapy, in the form of monoclonal antibodies, that clear Aβ. Interest in this approach is based on a disease model in which Aβ aggregation is posited as the disease trigger and other disease facets, including tau propagation, chronic neuroinflammation, brain atrophy, and ultimately diminished cognition all occur downstream.^3^ Anti-Aβ antibodies aim to mitigate such symptoms by clearing the putative trigger. Dozens of monoclonal antibodies (mAbs) targeting various species of Aβ have been developed and investigated in clinical trials.^4–8^ Though many have failed to reach their trial benchmarks, adcunumab (Aduhelm) was FDA approved via the Accelerated Approval Program in 2021 and lecanemab (Leqembi, 2023) and donanemab (Kisunla, 2024) have since received full FDA approval. These drugs are the first to be approved for AD in ∼20 years. However, the effects of these treatments on cognitive decline is small, and some have questioned whether the associated therapeutic benefits are clinically meangingful.^9^ This class of antibodies is also associated with side effects – most notably amyloid-related imaging abnormalities (ARIA), which is due to edema and/or hemorrhage in the brain.^10^ In the aducanumab (EMERGE and ENGAGE), lecanemab (Clarity AD), and donanemab (TRAILBLAZER-ALZ2) clinical trials, 12.6-36% of participants receiving drug experienced ARIA-Edema (compared to 0.8-3% of placebo group) and 17.3-35% experienced ARIA-Hemorrhage (compared to 7.2-10% of placebo group).^11–14^ While most episodes of ARIA resolved, a minority of ARIA cases have resulted in poor outcomes for patients, including a small number of fatalities.^10^ There is currently no therapy for AD that stops or reverses disease progression; those that slow cognitive decline do so at substantial risk to patients.

The next generation of experimental therapies for AD will likely need to address a broader range of factors beyond Aβ accumulation, and chronic inflammation has emerged as a potential major driver of disease. Microglia and astrocytes, both glial cells, become activated in response to Aβ and other features of the AD niche.^15,16^ Microglia take on a pro-inflammatory phenotype and produce inflammatory cytokines, including interleukin-1 (IL-1α), which induces synapse loss, and tumor necrosis factor (TNF), which can directly cause neuronal death.^17,18^ Significantly, these inflammatory signals from microglia induce pro-inflammatory reactive astrocytes.^19^ Reactive astrocytes are highly present in AD, are neurotoxic, and contribute to cerebrovascular dysfunction.^20–22^ New AD treatment approaches that mitigate neuroinflammation and that offer targeted support for CNS cells are needed and would represent a shift from the existing treatment paradigm requiring Aβ or tau protein clearance.

Engineered cell therapies, in which cells are programmed to survey their environment and exert therapeutic functions, have shown great success in other diseases – most notably chimeric antigen receptor T (CAR-T) cells in oncology. CAR-T cell therapy involves *ex vivo* transduction of a patient’s T cells to program them with an antibody-based receptor that can specifically recognize a tumor-associated antigen, thus coupling recognition of a tumor antigen to activation of the native T cell effector mechanisms.^23^ Six CD19 or B-cell maturation antigen (BCMA) CAR-T cell products are FDA-approved to treat blood cancers, including lymphomas, leukemias, and multiple myeloma.^24,25^ Building on the success of CAR-T therapy, CAR macrophages or microglia (CAR-M) have been investigated pre-clinically for AD; in these designs, the CAR is constructed from an anti-Aβ antibody fragment and results in macrophage activation through Fc receptor signaling, leading to phagocytosis of Aβ.^26,27^ Additionally, CAR-Astrocytes (CAR-A) have shown reduction in Aβ burden in 5xFAD mice and coordinate responses in astrocytes and microglia.^28^ While these approaches capitalize on native phagocytic function, the intended outcome still depends on Aβ clearance, which has not shown evidence of substantially modifying disease progression. To date, the success of engineered cell therapies has not been translated to neurodegenerative disease.

Here we present a potential cell-based therapy platform for AD, where cells can intelligently survey their microenvironment for disease markers (i.e., the Aβ plaque niche). As with CAR technology, this is accomplished through a synthetic receptor capable of recognizing Aβ. However, by utilizing the synthetic Notch (synNotch) platform,^29^ we design cells capable of responding to Aβ by activating a selected gene expression program; unlike CAR-M cells, the output of these cells is independent of Aβ clearance. SynNotch is based on the native juxtacrine Notch signaling pathway, in which recognition of the ligand (i.e., Delta and Jagged family members) triggers sequential cleavage by an ADAM metalloproteinase and γ-secretase within the transmembrane core of Notch, freeing the intracellular domain to translocate to the nucleus and drive transcription of downstream genes (Fig. 1A).^30^ To construct an Aβ-sensitive synNotch receptor, the extracellular and intracellular domains of Notch are exchanged for synthetic components: the extracellular domain for the single-chain variable fragment (scFv) of an anti-Aβ antibody, and the intracellular domain for a synthetic transcription factor (TF). The transmembrane core, where proteolysis occurs, is conserved in synNotch, allowing Aβ recognition to drive cleavage and release the TF, which translocates to the nucleus to drive expression of a programmed transgene (Fig. 1A). Cells that express these receptors sense Aβ as a marker of disease pathology and, via synNotch, respond with the expression of a programmed transgene. This Aβ-synNotch platform regulates expression of anti-inflammatory or neuroprotective transgenes. We use it in aged 5xFAD (named for the 5 Familial AD mutations; 3 in *APP*, 2 in *PSEN1*) transgenic mice, which show significant amyloid plaque burden and impaired memory and cognitive function by 6 to 9 months of age.^31,32^ Taken together, we demonstrate the capacity of the Aβ-synNotch system to engineer central nervous system (CNS) cells to overcome deleterious factors within the AD disease niche.

**Figure 1.**
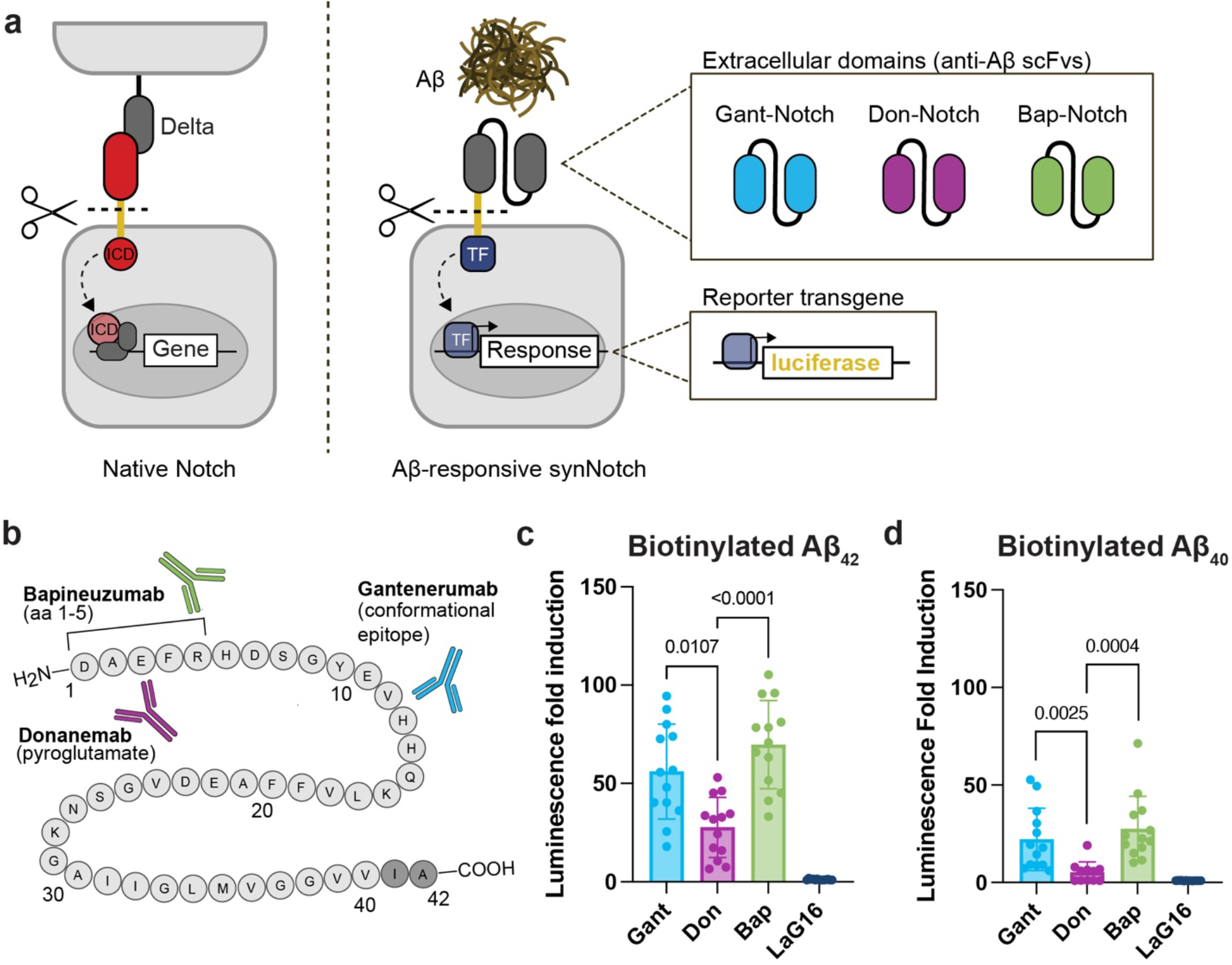
Engineered synNotch receptors enable programmable cellular recognition of immobilized, synthetic Aβ. (A) Schematic of Notch and synNotch. Notch recognizes its ligand Delta presented in *trans* by a neighboring cell, resulting in cleavage of the transmembrane core and releasing the intracellular domain to the nucleus for target gene regulation. Aβ-synNotch receptors were constructed by replacing the Notch1 extracellular domain with anti-Aβ monoclonal antibody single chain variable fragments (scFvs) and the intracellular domain (ICD) with the tet-transactivator synthetic transcription factor (TF). ScFvs were generated from anti-Aβ antibodies gantenerumab (Gant-Notch), donanemab (Don-Notch), and bapineuzumab (Bap-Notch). Here, activation of synNotch receptors by Aβ will result in expression of a luciferase transgene. (B) Depiction of gantenerumab, donanemab, and bapineuzumab epitopes on the Aβ peptide. (C) Aβ-driven luciferase transgene expression of L929 fibroblasts expressing Bap-Notch, Don-Notch, and Gant-Notch plated on biotinylated Aβ_42_ immobilized to plate surfaces via streptavidin. Data shown as fold induction of luminescence compared to cells plated on a control, Aβ-free surface. LaG16-Notch is a negative control (GFP-responsive) receptor. *n* = 13 from 3 independent experiments; mean±SD; Welch ANOVA with Dunnett’s T3 multiple comparisons. All groups are significantly different (p<0.05) from LaG16-Notch control (statistics not shown). (D) Fold induction of Aβ-driven luciferase transgene luminescence of synNotch cells plated on immobilized biotinylated Aβ_40_. *n* = 11-13 from 3 independent experiments; mean±SD; Welch ANOVA with Dunnett’s T3 multiple comparisons. All groups are significantly different from LaG16-Notch control unless shown.

## Results

### SynNotch receptors constructed from anti-Aβ monoclonal antibodies enable Aβ-driven transgene expression

Receptors were designed from a panel of anti-Aβ monoclonal antibodies previously tested in clinical trials for AD: gantenerumab (binds conformational epitope found in fibrillar Aβ);^5^ donanemab (recognizes N-terminal truncated pyroglutamate form of Aβ [Aβp3-42] found in plaques)^33^; bapineuzumab (recognizes amino acids 1-5; binds soluble and insoluble Aβ_40/42_)^34^ (Fig. 1B). These receptors were denoted as Gant-Notch, Don-Notch, and Bap-Notch, respectively. For initial characterization, the receptors were expressed under the EF1A promoter in mouse L929 fibroblasts by lentiviral transduction. Cells were sorted for matched levels of receptor expression by fluorescence-activated cell sorting (FACS) using a c-myc tag incorporated on the N-terminus of receptors (Supplementary Fig. 1A-B). Notably, prior to cell sorting, only a small fraction of fibroblasts successfully expressed Don-Notch, though receptor expression persisted in the sorted population (Supplementary Fig. 1C-D). Cells were programmed to express mCherry fluorescent protein and firefly luciferase in response to receptor recognition of Aβ. To characterize the ability of the synNotch receptors to recognize Aβ, we immobilized C-terminally biotinylated, synthetic Aβ_42_ and Aβ_40_ to streptavidin-coated well plates and seeded synNotch cells. Gant-Notch, Don-Notch, and Bap-Notch recognized synthetic Aβ_42_ immobilized to the plate surface, as indicated by Aβ-dependent luciferase transgene expression. Fold inductions of luminescence compared to synNotch cells plated on a control, Aβ-free surface range from 28x (Don-Notch) to 70x (Bap-Notch) (Fig. 1C). Gant-Notch and Bap-Notch also recognized Aβ_40_, the more prevalent but less pathogenic Aβ variant,^35,36^ immobilized by biotin-streptavidin interaction, indicated by luciferase transgene expression (Fig. 1D). This is consistent with the reported antibody epitopes, though overall, receptors exhibited a lower fold activation on Aβ_40_ than on Aβ_42_. Notably, Don-Notch, constructed from the plaque-specific antibody donanemab, does not recognize Aβ_40_ when immobilized via biotin-streptavidin interaction. SynNotch activation, as measured by luciferase activity, was not detected in cells expressing a control, GFP-responsive LaG16-Notch receptor, confirming that activation is dependent on selectively programming synNotch with Aβ-detection motifs.

We also constructed synNotch receptors with recognition domains derived from the other two anti-Aβ monoclonal antibodies approved by the FDA for AD: aducanumab (recognizes amino acids 3-7; binds soluble oligomers and insoluble fibrils of Aβ_42_) (Ad-Notch);^37,38^ and lecanemab (binds a conformational epitope found in Aβ protofibrils) (Lec-Notch)^39^ (Supplementary Fig. 2A). Ad-Notch and Lec-Notch did not recognize synthetic Aβ in our studies (Supplementary Fig. 2B-C). To validate that both Lec- and Ad-Notch receptors are functional in the fibroblasts, we treated the cell cultures with 1 µm beads decorated with antibodies specific to the c-myc epitope tag. Such beads serve as surrogates for Aβ ligand, as the N-terminal domain of the synNotch receptor contains the c-myc tag and thus engages the beads in a manner similar to immobilized ligand (i.e., Aβ). Results reveal that the Ad-Notch and Lec-Notch receptors can be activated (Supplementary Fig. 2B-C), suggesting that the lack of response to Aβ is not due to a defect in the receptor design. A previous report of an Aβ-specific receptor constructed from aducanumab does recognize Aβ; this may be due to differences in construct design, including orientation of the variable domains of the scFv, or due to differences in Aβ preparation^40^ Due to their underperformance in converting Aβ presence to a productive, engineered cell response, Lec-Notch and Ad-Notch were not included in subsequent experiments.

### Aβ-synNotch receptors differentially recognize Aβ_42_ and Aβ_40_

To determine whether the Aβ-synNotch receptors recognize unmodified Aβ (i.e., not biotinylated), coated the culture surface with synthetic Aβ_42_ via passive adsorption. Gant-Notch, Don-Notch, and Bap-Notch recognized synthetic Aβ_42_ adsorbed to the plate, as demonstrated by the fold induction in luciferase transgene expression compared to cells plated on a control surface (Fig. 2A). Remarkably, for all three receptor variants, passive adsorption of unmodified Aβ_42_ results in higher luciferase fold induction than on biotinylated Aβ_42_, ranging from 53x (Don-Notch) to 155x (Bap-Notch). We were also interested in whether the synNotch receptors recognize Aβ supplemented in the medium. Gant-Notch, Don-Notch, and Bap-Notch recognize Aβ_42_ in the culture medium, though Bap-Notch resulted in significantly higher luminescence fold induction than all other receptors (Fig. 2B). Of note, synNotch activation requires a mechanical force of 4-12 pN to activate; this cannot be provided by soluble, monomeric ligands, and thus synNotch generally requires an immobilized ligand.^29^ We have previously shown that oligomeric ligands like TGFβ-1, VEGF, TNF, and a dimeric form of GFP can activate cognate synNotch receptors.^41^ Accordingly, we hypothesized that the oligomeric nature of Aβ (specifically, multiple instances of the same epitope on an Aβ oligomer) may similarly allow for recognition of Aβ_42_ supplemented to the medium. Unlike Aβ_42_, Aβ_40_ must be anchored to the surface to result in synNotch activation (Fig. 2C); medium supplemented Aβ_40_ resulted in less than 2-fold induction in synNotch activation (Fig. 2D). This discrepancy between Aβ_42_- and Aβ_40_-driven synNotch activation is likely due to the decreased propensity of Aβ_40_ to form stable aggregates compared to Aβ_42_, resulting in fewer epitopes available for multiple synNotch receptors to simultaneously engage Aβ_40_ to generate the mechanical stress required for activation.^42^ After screening the panel of receptors for activation on a variety of Aβ ligands, we selected Bap-Notch for subsequent work due to its high sensitivity in detecting immobilized Aβ_42_ and Aβ_40_, as well as Aβ_42_ supplemented to the medium, where it resulted in a significantly higher fold induction in luciferase transgene expression than either Gant-Notch or Don-Notch.

**Figure 2.**
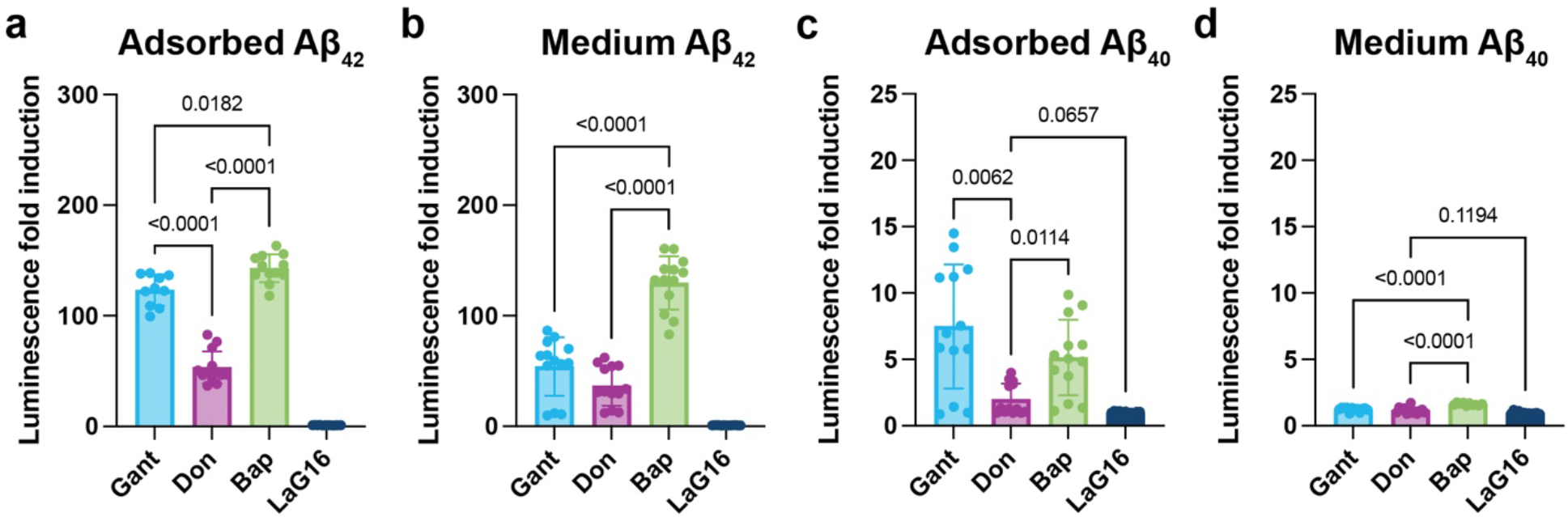
Aβ-synNotch receptors display differential levels of recognition of unmodified Aβ peptides. Fold induction of Aβ-driven luciferase transgene luminescence of synNotch cells plated on (A) Aβ_42_ adsorbed to the plate, (B) Aβ_42_ supplemented to the medium, (C) Aβ_40_ adsorbed to the plate, and (D) Aβ_40_ supplemented to the medium, compared to cells plated on a control, Aβ-free surface. LaG16-Notch is a negative control (GFP-responsive) receptor. *n* = 10-13 from 3 independent experiments, with outliers removed; mean±SD; Welch ANOVA with Dunnett’s T3 multiple comparisons. All groups are significantly different from LaG16-Notch control unless shown.

### Aβ-synNotch governs therapeutic transgene expression in human glia

We next sought to express Bap-Notch in microglia, the resident macrophages of the brain, which maintain brain health through phagocytosis of debris and promotion of inflammation during a pathological challenge.^43–45^ While crucial to homeostatic brain health, these functions become dysregulated or chronically activated in AD,^16,46^ making microglia a key target for reprogramming via synNotch circuits. Additionally, microglia replacement therapy is currently being investigated for diseases involving dysfunctional microglia, most notably with microglial replacement by bone marrow transplantation in patients with adult-onset leukoencephalopathy with axonal spheroids and pigmented glia (ALSP).^47,48^ While microglia replacement therapy for AD is still in a preclinical stage, its success in mouse models supports interest in a microglia-based therapeutic strategy for AD.^49–51^ Here, immortalized human microglia were engineered to express the Bap-Notch receptor via lentiviral spinoculation. Bap-Notch microglia recognize synthetic Aβ_42_, both immobilized to the cell culture surface via streptavidin-biotin interaction and supplemented to the medium, as compared to microglia expressing the off-target LaG16-Notch receptor, indicated by luciferase transgene expression (Fig. 3A-B).

**Figure 3.**
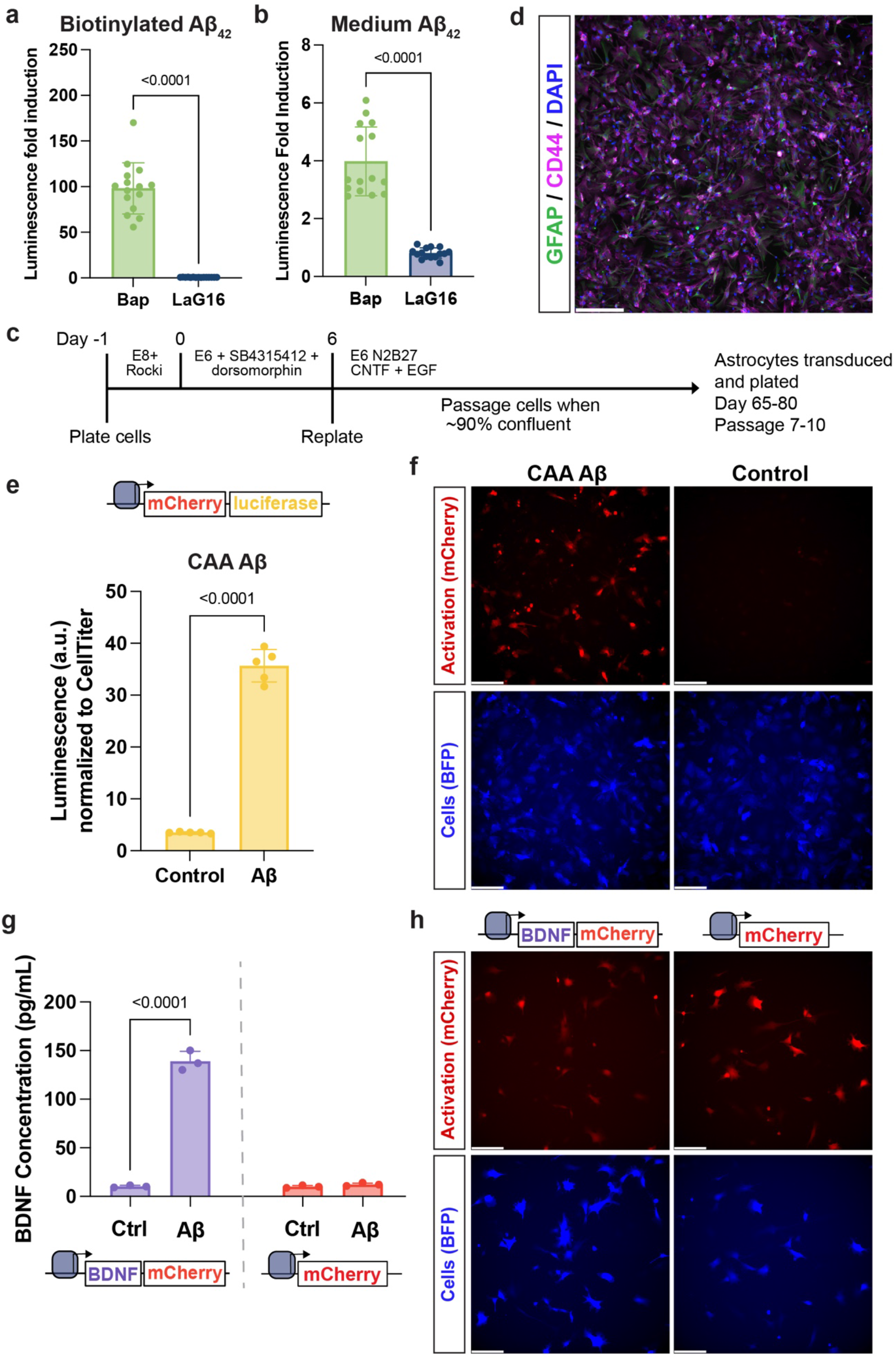
SynNotch glia regulate programmable Aβ-dependent transgene expression. Fold induction of Aβ-driven luciferase transgene luminescence of Bap-Notch microglia plated on (A) biotinylated Aβ_42_ immobilized to well surfaces via streptavidin and (B) Aβ_42_ supplemented to the medium compared to cells plated on control surface. *n* = 15 from 3 independent experiments; mean±SD; Welch’s *t-*test. (C) Schematic of astrocyte derivation from hiPSCs. (D) Expression of astrocytic markers CD44 and GFAP. DAPI shows cell nuclei. Scale bar = 200 µm. (E) Aβ-driven firefly luciferase luminescence from Bap-Notch astrocytes plated on Aβ isolated from CAA patients. Luminescence is normalized to CellTiter Fluor. *n* = 5; mean±SD; Welch’s *t-*test. (F) Representative Aβ-dependent mCherry expression in Bap-Notch astrocytes plated on CAA-Aβ. Scale bar = 200 µm. (G) Aβ-dependent BDNF expression (measured by ELISA) in Bap-Notch astrocytes programmed with either a BDNF-mCherry transgene or an mCherry only transgene and plated on immobilized synthetic Aβ_42_. *n* = 3; mean±SD; one-way ANOVA with Tukey’s multiple comparisons. (H) Representative Aβ-dependent mCherry expression in Bap-Notch astrocytes programmed with either a BDNF-mCherry transgene or an mCherry only transgene and plated on immobilized synthetic Aβ_42_. Scale bar = 200 µm.

We additionally expressed Bap-Notch in astrocytes, glial cells that have a major role in maintaining the blood-brain barrier (BBB), regulating synapses, and coordinating the immune response in the brain.^52^ In AD, these roles become dysregulated.^53,54^ In particular, astrocytes deeply interact with and shape the amyloidogenic microenvironment by phagocytosing Aβ and releasing pro-inflammatory factors.^15,20^ Our goal is to capitalize on the native homeostatic function of astrocytes by enhancing Aβ recognition and implementing programmable therapeutic responses. Astrocytes were differentiated from CC3 human iPSCs via dual-SMAD inhibition (SB431542 and dorsomorphin) followed by culture in the presence of EGF and CNTF (Fig. 3C-D, Supplementary Fig. 3).^21^ To encourage expression of the receptor in astrocytes, the constitutive promoter driving the synNotch receptor was transitioned from EF1A to an astrocyte-specific truncated glial fibrillary acidic protein (GFAP) promoter (gfaABC_1_D).^55^ Astrocytes were engineered to express the Bap-Notch receptor via lentiviral transduction. We were interested in whether the synNotch receptors could recognize patient-derived Aβ, in addition to synthetic Aβ. Aβ isolated from brain bank donors with cerebral amyloid angiopathy (CAA) was added to the culture medium of astrocytes. Bap-Notch astrocytes recognized the patient-derived Aβ, as indicated by the Aβ-driven upregulation of firefly luciferase; luminescence is normalized to CellTiter fluorescence to account for differences in astrocyte number (Fig. 3E). The same astrocytes also upregulate an mCherry transgene in response to the patient-derived Aβ (Fig. 3F). These results demonstrate that human brain cells engineered with Aβ-synNotch receptors recognize human brain-derived Aβ and respond with programmable transgene output.

We were next interested in using synNotch astrocytes to regulate potentially therapeutic gene expression programs in response to Aβ. Brain-derived neurotrophic factor (BDNF) is a neuronal growth factor that has shown great promise in encouraging neuronal growth and synapse formation after brain injury in non-human primates (NHPs).^56,57^ It has been investigated as a therapy for neurodegeneration when delivered in AAV, however, widespread delivery of BDNF to the brain has been associated with severe side effects (weight loss, sensory disturbances, and inappropriate cellular migratory patterns).^58^ To avoid these, more recent strategies have demonstrated success with delivery of BDNF-AAV via direct injection to the entorhinal cortex in non-human primates.^59^ When used as a synNotch response transgene, ectopic BDNF expression is regulated by local Aβ accumulation, thus allowing for specificity in delivery. To explore the feasibility of using synNotch to govern BDNF expression, Bap-Notch astrocytes were engineered to express BDNF and mCherry in response to Aβ and were plated on synthetic Aβ_42_. Bap-Notch astrocytes expressed BDNF only when cultured with Aβ (Fig. 3G). Astrocytes engineered to express only mCherry as a synNotch transgene (*i.e.* no BDNF) implement a synNotch-governed response to Aβ, as indicated by mCherry expression (Fig. 3H); however, such cells did not upregulate BDNF production, confirming that enhanced BDNF expression is potentiated by synNotch and is not attributable to a native astrocyte response.

### Bap-Notch attenuates astrocyte inflammation in response to Aβ

Chronic neuroinflammation, mediated by astrocytes and microglia, plays a major role in AD progression. Many AD risk genes (e.g., *APOE, TREM2, CD33*) identified in genome-wide association studies (GWAS) skew immune cell function toward a pro-inflammatory phenotype.^60–62^ Additionally, elevated levels of inflammatory cytokines are found in the cerebral spinal fluid (CSF) of AD patients,^63^ suggesting the AD brain may be conducive to a reactive astrocyte phenotype. Indeed, pro-inflammatory, reactive astrocytes have been identified in post-mortem brain tissue in AD.^20^ Further, *in vitro* models suggest that microglial responses to amyloid produced by neurons expressing APP with a Swedish mutation (APP^Swe^) exacerbate astrocyte reactivity.^64^ We were therefore interested in whether we could use synNotch to re-route this inflammatory, reactive astrocyte phenotype via an engineered response to Aβ. To this end, we programmed Bap-Notch astrocytes to mitigate pro-inflammatory cues via expression of soluble TNF receptor (sTNFR1) and IL-1 receptor antagonist (IL-1Ra) in response to Aβ. A control astrocyte line was generated that expressed SEAP downstream of synNotch. Bap-Notch astrocytes were plated on synthetic Aβ_42_ for 48 hr prior to supplementing the medium with IL-1α (5 ng/mL) and TNF (10 ng/mL) to induce a reactive astrocyte phenotype (Fig. 4A).^20,64–66^ After 48 hr exposure to cytokines, Bap-Notch astrocytes engineered to antagonize TNF and IL-1α dramatically upregulated sTNFR1 and IL-1Ra in an Aβ-dependent manner (approximately 40 ng/mL) (Fig. 4B-C). Correspondingly, qRT-PCR revealed that Bap-Notch mediated significant attenuation of reactive genes interleukin-6 (*IL6)*, complement C3 (*C3*), and granulocyte-macrophage colony stimulating factor (*CSF*2) expression when Aβ was present to drive expression of the anti-inflammatory transgenes (Fig. 4D). Bap-Notch SEAP-astrocytes, however, did not mitigate reactive gene expression in response to Aβ (Fig. 4E), despite responding potently to Aβ, as represented by significant Aβ-driven SEAP production (Supplementary Fig. 4D). These results reflect the ability to deliberately modulate cell responses based on transgene selection and demonstrate the utility of cytokine antagonist transgenes in moderating the inflammatory environment that leads to glial reactivity and eventual neuron loss.

**Figure 4.**
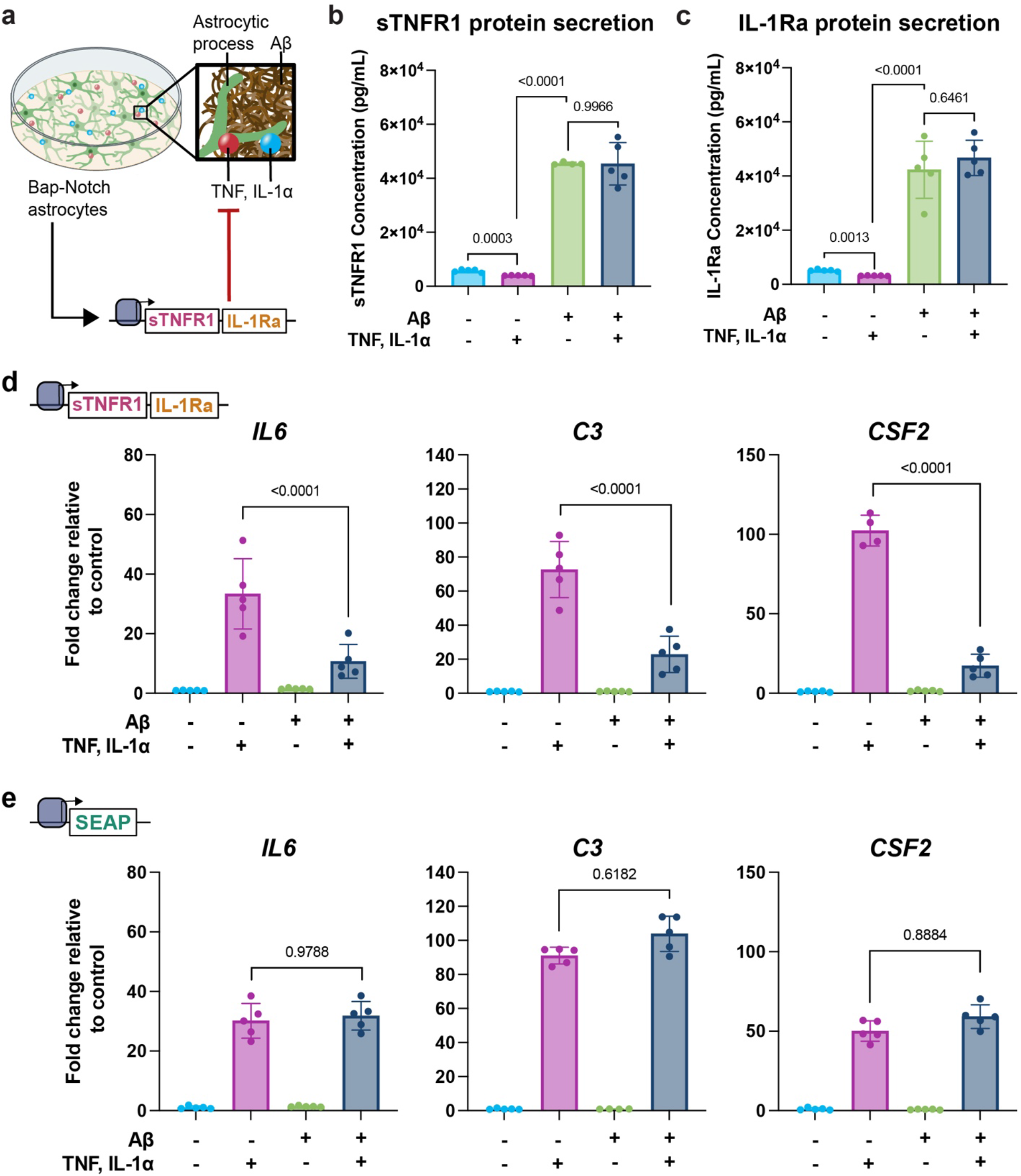
Bap-Notch attenuates astrocyte reactivity. (A) Schematic of experimental configuration. Bap-Notch astrocytes are plated on synthetic Aβ_42_ embedded in Matrigel. Inflammatory cytokines TNF and IL-1α are added to the medium. Recognition of Aβ results in the astrocytes expressing inflammatory cytokine antagonists sTNFR1 and IL-1Ra. sTNFR1 (B) and IL-1Ra (C) transgene expression (measured by ELISA) of Bap-Notch astrocytes plated on immobilized Aβ_42_ and treated with TNF (10 ng/mL) and IL-1α (5 ng/mL). *n* = 4-5, with outliers removed; mean±SD; two-way ANOVA with Tukey’s multiple comparisons. (D) Reactive gene expression in Bap-Notch astrocytes that express an sTNFR1 and IL-1Ra transgene in response to Aβ. Bap-Notch astrocytes were plated on either an Aβ or control surface, and half were treated with TNF and IL-1α. *n* = 4-5, with outliers removed; mean±SD; two-way ANOVA with Tukey’s multiple comparisons. All groups treated with IL-1α and TNF are statistically different from groups without cytokines (differences not shown). (E) Reactive gene expression in Bap-Notch astrocytes that express a SEAP reporter in response to Aβ. *n* = 4-5 due to outlier removal; mean±SD; two-way ANOVA with Tukey’s multiple comparisons. All groups treated with IL-1α and TNF are statistically different from groups without cytokines (differences not shown).

### Aβ-responsive synNotch receptors detect Αβ in the 5xFAD transgenic mouse model

We sought to demonstrate the feasibility and functional consequence of delivering Aβ-synNotch cells in a model of transplanted cell therapy. We expressed Bap-Notch in mouse mesenchymal stromal cells (MSCs) via lentiviral transduction and sorted for a receptor-positive population via FACS. MSCs were chosen for this application for several reasons: (1) they are easily engineered to express synNotch receptors by lentiviral transduction and can be sorted to obtain a population of synNotch-expressing cells;^67–69^ (2) they can be transplanted to mice directly; and (3) they have been investigated as cell therapy for AD.^70–72^ To determine whether the Aβ-synNotch receptors can recognize Aβ *in situ*, we seeded Bap-Notch MSCs on brain organotypic slice cultures (OSCs) prepared from aged (11-month) hemizygous 5xFAD mice and wild-type (WT) controls (Fig. 5A). For each 5xFAD and WT mouse, half the total number of slices were seeded with Bap-Notch cells; the other half were seeded with GFP-responsive LaG16-Notch cells as a negative control. All synNotch MSCs were engineered to express SEAP and mCherry upon ligand engagement (*i.e.*, Aβ, GFP). We observed a significant increase in SEAP expression only in Bap-Notch cells seeded on slices from hemizygous 5xFAD mice (Fig. 5B). Levels of SEAP from Bap-Notch cells on WT OSCs were comparable to those measured from negative control LaG16-Notch cells on 5xFAD or WT OSCs, or from OSCs that did not receive MSC transplantation. To validate the SEAP findings, we cryosectioned OSCs and immunolabeled for Aβ and mCherry. The transplanted cells were visualized with constitutive BFP. Bap-Notch MSCs plated on the Aβ-containing hemizygous 5xFAD slices demonstrated strong mCherry positivity co-localized with Aβ deposits (Fig. 5C). As expected, wild-type tissue lacks Αβ, and transplanted MSCs (BFP+) are not mCherry+ (Fig. 5D). These results indicate that Bap-Notch cells implement potent responses to Aβ in native 5xFAD brain tissue.

**Figure 5.**
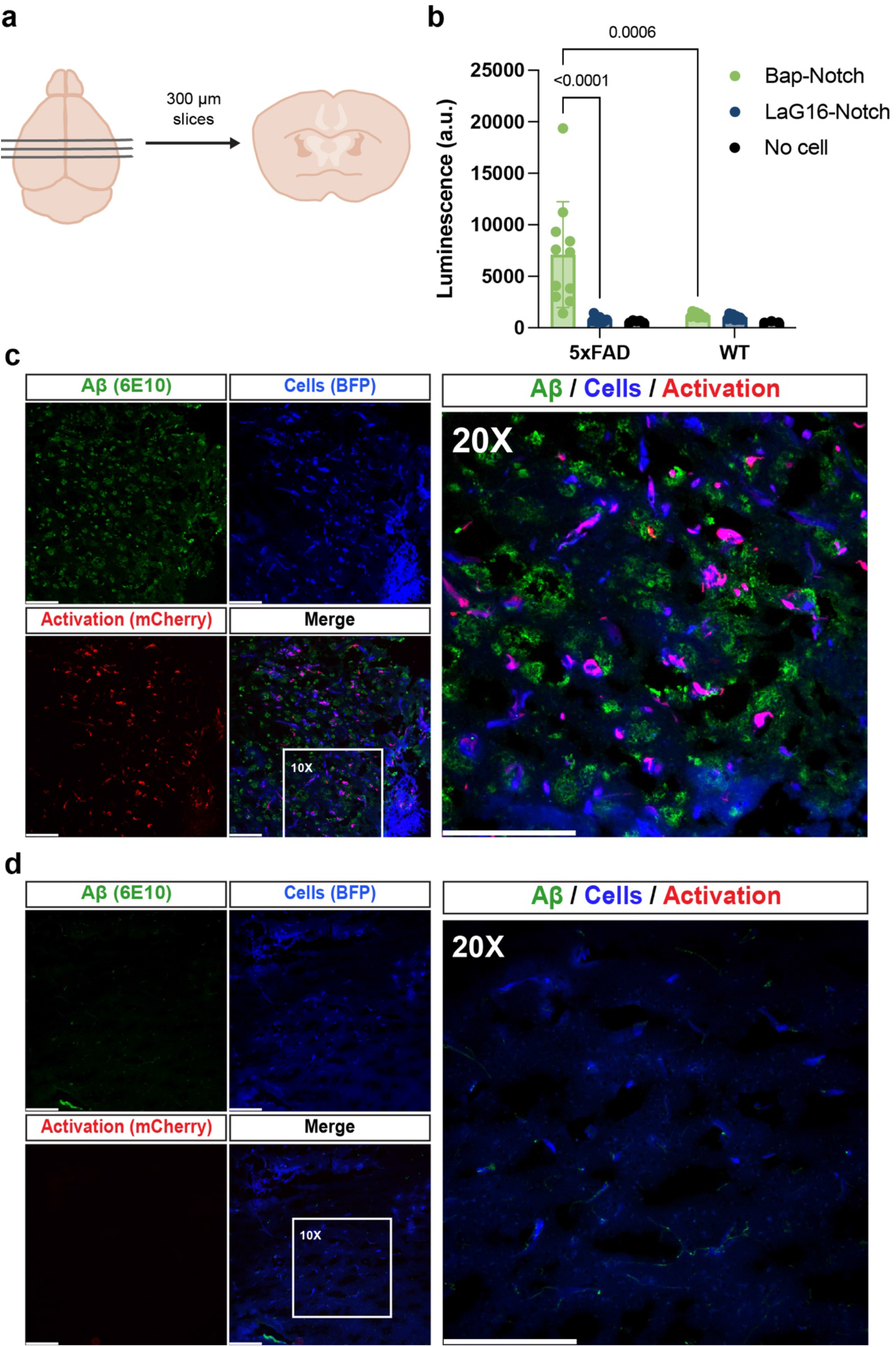
Recognition of amyloid *in situ* in organotypic slice cultures. (A) Schematic of coronal organotypic slice culture (OSC) preparation. (B) Aβ-dependent SEAP expression of Bap-Notch or LaG16-Notch MSCs plated on hemizygous 5xFAD or wild-type (WT) OSCs. *n* = 3-11 replicates; mean±SD; Welch ANOVA with Dunnett’s T3 multiple comparisons. (C-E) Fluorescence microscopy of Aβ (6E10 immunolabeled), constitutive BFP, and synNotch-driven mCherry (immunolabeled) in cryosections of hemizygous 5xFAD (C) or wild-type OSCs (D) treated with Bap-Notch MSCs. Scale bar = 200 µm.

Following confirmation that Bap-Notch recognizes Aβ deposited in the 5xFAD mouse brain, we delivered Bap-Notch MSCs directly to the brain of three aged (∼12-months-old) 5xFAD mice and three age-matched WT mice via stereotactic injection (Fig. 6A). As above, Bap-Notch MSCs were engineered to express SEAP and mCherry upon Aβ engagement. Three days after injection, cerebrospinal fluid (CSF) and blood serum were collected and brains were harvested and cryosectioned. We observed an increase in SEAP expression in the serum only in 5xFAD mice injected with Bap-Notch cells, compared to both WT cells treated with Bap-Notch cells and uninjected cohorts of 5xFAD and WT mice, indicating SEAP expression driven by Aβ recognition (Fig. 6B). Interestingly, we did not see an increase in SEAP in the CSF of 5xFAD mice (Fig. 6C); however, known reductions in brain volume in 5xFAD mice would be expected to result in ventricular enlargement and increased total CSF volume, resulting in further dilution of Aβ-driven SEAP in 5xFAD.^73,74^ We probed for activated cells via immunofluorescence in cryosections of injected brains. Bap-Notch cells constitutively expressed GFP, allowing visualization of deposited cells in the brain. GFP+ cells largely deposited together, with some migration from the injection site along the corpus callosum. Additionally, some cells remained in the injection tract through the cortex at this timepoint (Supplementary Fig. 5A). The pattern of cell deposition was similar in 5xFAD and wild-type mice and Fig. 6D-E depict cells in the same region (Supplementary Fig. 5A). As expected, 5xFAD mice had significant Aβ deposition throughout the cortex and hippocampus, stained by the 6E10 antibody (Fig. 6D, Supplementary Fig. 5B). Only cells in 5xFAD mice showed mCherry expression, driven by Bap-Notch recognition of Aβ (Fig. 6D-E). Together, these data demonstrated that Bap-Notch cells delivered directly to the brains of aged mice recognize Aβ and subsequently express programmed transgenes.

**Figure 6.**
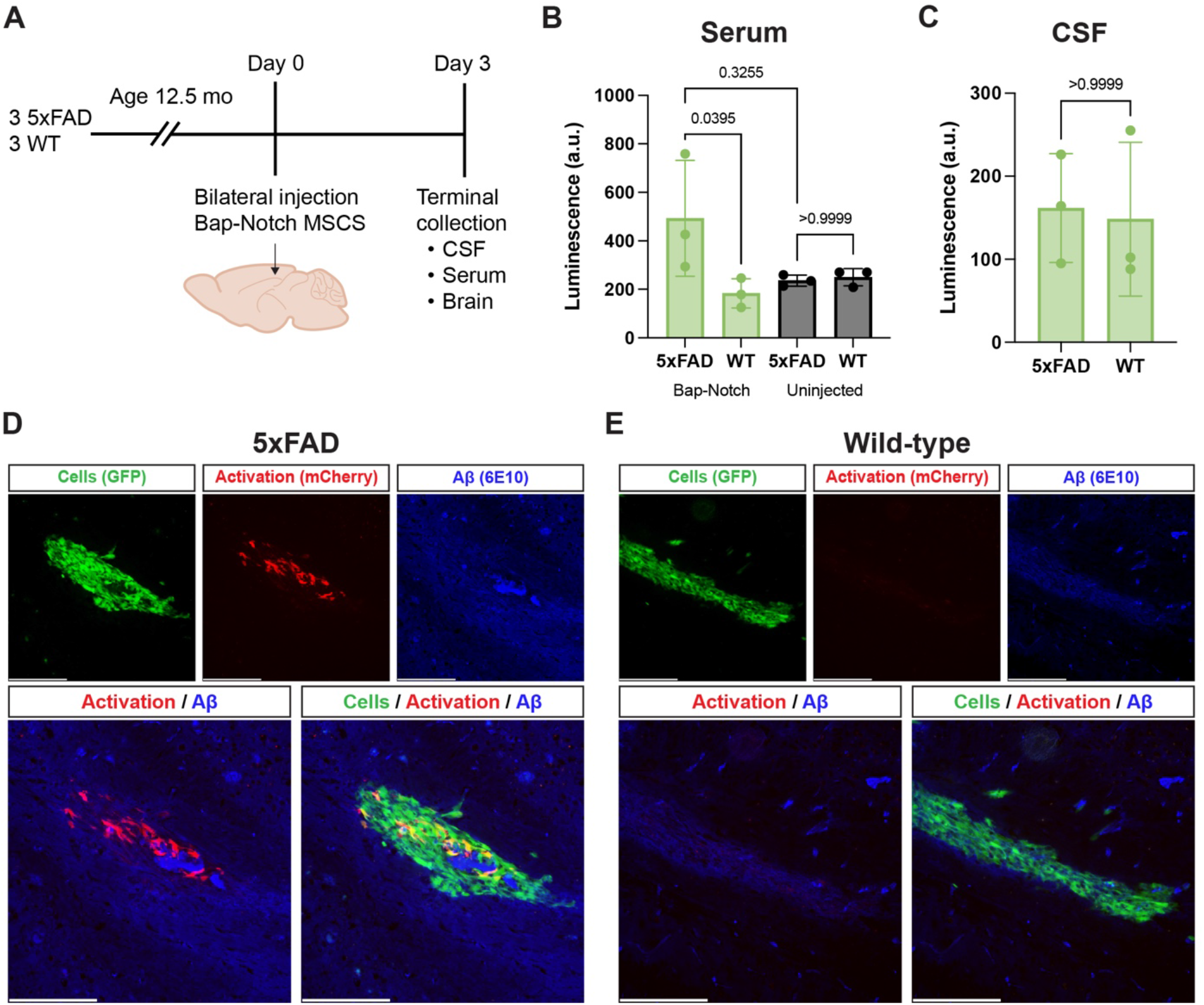
Aβ-driven activation of Bap-Notch MSCs transplanted to the brains of aged 5xFAD mice. (A) Schematic of experiment. Three hemizygous 5xFAD and 3 wild-type mice were aged for 12 months. 150,000 Bap-Notch MSCs were injected bilaterally. After 72 hr, CSF and blood serum were harvested for analysis by SEAP activity assay and brains were harvested for cryosectioning. (B-C) Αβ-dependent SEAP expression in the serum (B) or CSF (C) collected from Bap-Notch MSCs transplanted to 12-month-old 5xFAD or wild-type mice. *n=*3 mice; mean±SD; Kruskal-Wallis test (B), Mann-Whitney test (C). (D-E) Fluorescence microscopy of transplanted cells (constitutive GFP, immunolabeled), Aβ (6E10 immunolabeled), and synNotch-driven mCherry (immunolabeled) in cryosections of 12-month-old 5xFAD (D) or wild-type (E) mice with Bap-Notch MSCs transplanted to the cortex. Scale bar = 200 µm.

### Aβ-synNotch platform is compatible with CNS-targeting gene delivery vehicles

While transplanted, engineered cells represent one potential avenue for deploying synNotch cells in AD therapy, transducing endogenous astrocytes to express the synNotch circuits offers an alternative translational approach that may be more compatible with clinical utility as an off-the-shelf gene therapy. Towards this goal, we constructed an AAV plasmid capable of delivering the Aβ-detecting receptor and response transgenes simultaneously. The packaging capacity of AAV is capped at 4.7 kb of DNA.^75^ Expression of the complete synNotch circuit would therefore require dual transduction, which may drastically limit the number of cells able to drive therapeutic output in response to Aβ. The Synthetic Intramembrane Proteolysis Receptors (SNIPR) platform is a derivative of synNotch, with one common variant composed of truncated Notch transmembrane and juxtamembrane domains.^76,77^ The smaller size of such SNIPR architectures allows for expression of both the receptor and the response transgenes from one AAV vector. Thus, we constructed a Bap-SNIPR receptor. To validate Bap-SNIPR responsiveness to Aβ, Bap-SNIPR and Bap-Notch L929 fibroblast cells were plated on immobilized Aβ_42_ and Aβ_40_. Bap-Notch and Bap-SNIPR resulted in similar fold changes in luciferase expression compared to control, Αβ-free conditions (Supplementary Fig. 6A-B). Bap-SNIPR and Bap-Notch also display comparable recognition of unmodified Aβ_42_ and Aβ_40_ when these species are supplemented in the medium or adsorbed to the plate surface (Supplementary Fig. 6C-E). These results demonstrate the feasibility of installing the artificial Aβ-responsive signaling channel in a format compatible with AAV, which are therapeutically relevant and widely investigated for use as gene delivery vehicles in the CNS.

Delivery of synNotch circuits as a gene therapy will result in the synNotch cell retaining patient mutations; of particular concern are mutations in the *PSEN1* gene which result in familial Alzheimer’s disease (FAD). *PSEN1* encodes for presenilin 1, a component of the γ-secretase complex, which functions in Aβ processing from the amyloid precursor protein (APP), but is also crucial to the function of the synNotch and SNIPR platforms.^29,76^ These cases constitute only an extremely small percentage of AD cases: while *PSEN1* mutations are the most common driver of autosomal dominant early onset AD (EOAD) (43%), these cases account for less than 1% of AD cases overall.^78–80^ However, it would be beneficial to understand whether our Aβ-detecting receptors function in cells with disease-causing mutations in *PSEN1.* We expressed Bap-Notch in patient-derived fibroblasts (Coriell Institute) with mutations known to affect Aβ processing from APP, including a 56-year-old man and a 56-year-old woman with cognitive decline. We also expressed Bap-Notch in an age-matched control fibroblast line with no known disease, confirmed by brain MRI scan and neurological exam. Aβ recognition will result in the expression of firefly luciferase and mCherry. Bap-Notch functioned in all three donors, including those with *PSEN1* mutations that may affect γ-secretase processing, as indicated by Aβ-driven expression of luciferase and mCherry (Supplementary Fig. 7). This lends confidence to the use of Aβ-detecting synNotch circuits as a gene therapy, including potentially in cases of *PSEN1* FAD.

## Discussion

Here, we present a new cell therapy approach for AD. In the last several decades, many monoclonal antibodies targeting Aβ have been developed as passive immunotherapy for AD treatment.^37,81^ We repurposed these antibodies to generate a panel of Aβ-sensitive synNotch receptors. We use these receptors to program cells that recognize Aβ as a marker of local disease and consequently execute conditional, prescribed functions independent of Aβ clearance. Here, we demonstrate that synNotch receptors constructed from anti-Aβ mAbs with different epitopes allow for differential recognition between distinct species of Aβ presented in varied contexts (i.e., immobilized or medium-supplemented). We express Bap-Notch, the receptor constructed from bapineuzumab, in glial cells, including human immortalized microglia and iPSC-derived astrocytes, and use it to express potential therapeutic transgenes in an Aβ-dependent manner. The expression of the anti-inflammatory transgenes IL-1Ra and sTNFR1 in response to Aβ attenuates a reactive astrocyte gene profile. We also implement the Aβ-synNotch system in MSCs, the cell type that makes up Lomecel-B, an investigational cell therapy for AD. ^71,72^ We deliver Bap-Notch mMSCs directly to the brain of aged 5xFAD mice and show that they can recognize Aβ *in vivo*, based on the expression of response transgenes SEAP and mCherry. Together, these results illustrate that the Aβ-synNotch platform serves as a potential engineered therapy for AD.

The modularity of synNotch lends immense utility to this system for developing targeted therapies for AD. Here, we have used the platform to express both neuroprotective (e.g., BDNF) and anti-inflammatory (e.g., sTNFR1, IL-1Ra) transgenes from the same Aβ-synNotch receptor. SynNotch-astrocytes regulated BDNF production in an Aβ-dependent manner. Additionally, Aβ-dependent sTNFR1 and IL-1Ra expression by Bap-Notch astrocytes significantly attenuated astrocyte reactivity based on the expression of representative genes *IL6, C3,* and *CSF2.* These dramatic changes in inflammatory gene expression would be expected to influence the function of astrocytes, including on their canonical function of BBB maintenance. Future work will investigate whether anti-inflammatory synNotch astrocytes are able to preserve BBB integrity against inflammation-mediated disruption. This would be extremely relevant to AD, where accumulation of Aβ and chronic inflammation are frequently accompanied by vascular changes leading to BBB disruption.^84,85^

The use of the Aβ-synNotch system is not restricted to the therapeutic transgenes selected here. As more about the disease becomes understood, the platform can be adapted to regulate relevant therapeutic candidate transgenes. Because synNotch transcriptional responses are potentiated by direct contact with immobilized ligand, the platform allows for the possibility of using synNotch to express therapeutic compounds that would be detrimental if delivered systemically to the brain, including BDNF, and instead gates their production on disease-dependent features that accumulate as AD progresses.

Translation of the Aβ-synNotch system into preclinical models of AD is imperative to expand its utility in the context of an AD cell therapy. We first demonstrated the ability of Aβ-synNotch receptors to recognize Aβ *in situ* by delivering Bap-Notch MSCs to OSCs prepared from aged 5xFAD mice. This result allowed us to deliver Bap-Notch to living aged 5xFAD mice. Bap-Notch cells transplanted to the brains of 5xFAD mice demonstrated Aβ-dependent activation. MSCs are an excellent candidate for this sort of engineering, as they have been investigated in Phase I/II clinical trials for AD; in particular, one investigational AD therapy, Lomecel-B, is composed of allogenic MSCs.^71,72^ The delivery of *ex vivo* engineered Bap-Notch MSCs directly to the brains of mice mimics the deployment of CAR-T cell therapy, in which T cells are engineered *ex vivo* by lentivirus or γ-retrovirus before delivery.^86^ T cells engineered *ex vivo* with a synNotch regulating CAR expression, which allows additional layers of logic to be incorporated into the T cell design, are now being deployed in human clinical trials in the context of glioblastoma (NCT06186401).^82,83^ In the future, an *ex vivo* engineering strategy to deliver synNotch machinery may also be compatible with microglia replacement strategies, which could be done with synNotch HSCs by bone marrow transplant.^48^ Altogether, these considerations foreshadow widespread applications of an *ex vivo* engineering strategy that is compatible with use of allogenic cell sources.

An alternative to *ex vivo* cell engineering is directly engineering endogenous astrocytes, thus capitalizing on the native surveillance functions of mature CNS cells. Delivery of the synNotch circuit via CNS-targeted AAV may allow for delivery via systemic injection and could open the door to an off-the-shelf therapeutic that does not require *ex vivo* cell engineering. The AAV9 serotype intrinsically has the capability to cross the BBB and has been investigated in clinical trials for a variety of indications.^87–90^ However, the AAV9 transduction efficiency in the adult human brain is limited, motivating efforts to develop a capsid that exploits receptor-mediated transcytosis pathways, such as the human transferrin receptor (TfR1), to enhance brain delivery.^91^ The development of the PHP.eB capsid, which crosses the mouse BBB via the LY6A receptor and efficiently transduces murine brain cells, will allow this strategy to be first investigated in transgenic mouse models.^92^ Due to inherent AAV packaging capacity limitations (<4.7 kb), the construction and validation of the smaller Bap-SNIPR receptor, possessing the same Aβ recognition motif and similar intramembrane proteolysis regulation as the Bap-Notch receptor, represents a major step towards porting the programmable Aβ-responsive receptor system to AAV. Additionally, we have demonstrated that our receptors are still capable of detecting Aβ when expressing in cells with a genetic background that includes FAD-causing *PSEN1* mutations. Future work will deliver AAV to aged 5xFAD mice and confirm Aβ-dependent transgene expression *in vivo*.

Bap-Notch was selected for our work generating Aβ-dependent BDNF expression, attenuating astrocyte inflammation, and detecting Aβ in the 5xFAD mouse brain. Bap-Notch consistently resulted in a wide dynamic signaling range against Aβ in a variety of contexts and is expressed efficiently in multiple cell types, including fibroblasts, MSCs, microglia, and astrocytes. Notably, bapineuzumab, the antibody from which the Bap-Notch recognition domain is derived, can detect a range of Aβ species, including both soluble and insoluble Aβ species.^34,93^ This supports the case for using Bap-Notch to generate a synNotch-based cell therapy for AD. However, the other receptors constructed here, Gant-Notch and Don-Notch, also recognize both synthetic Aβ_42_ and Aβ_40_. It is interesting that there are differences in their ability to detect Aβ preparations, which is likely due to the different species of Aβ the antibodies are designed to detect. Future work characterizing the ability of the synNotch receptors constructed from the antibodies to recognize particular species may result in a cell capable of finely discriminating between Aβ species, as well as between Aβ and non-Aβ components of the periplaque niche. This may be of particular interest in the development of an Aβ-aggregation sensor system. Tau propagation sensors have been developed,^94^ but to our knowledge, there is no equivalent Aβ system.

In recent years, work by other groups has also begun to investigate glial engineering approaches for treating AD – to date, all have used engineered cellular responses to target Aβ for clearance. CARs, also constructed from previously developed anti-Aβ antibodies including aducanumab, have been expressed in both microglia and astrocytes.^27,28^ Both CAR-M and CAR-A increase phagocytosis of Aβ; CAR-A has been shown to result in Aβ clearance in aged 5xFAD mice. Another microglia engineering approach has been accomplished by encoding the Aβ-degrading enzyme neprilysin under the CD9 promoter (activated in microglia when proximal to Aβ deposits) by CRISPR gene editing.^51^ There has been a prior report of an Aβ synNotch receptor constructed from aducanumab.^40^ Notably, in this report, synNotch activation results in the expression of secreted chimeric anti-Aβ antibodies aducanumab and lecanemab, with the goal again being Aβ clearance. This Aβ synNotch system was expressed only in 3T3 fibroblasts, rather than in glia. SynNotch receptors targeting neuronal signal ganglioside (Gtb1) and apoptotic signal phosphatidylserine (PS) have been previously expressed in primary neonatal mouse astrocytes for detection of neurodegeneration.^95^ Here, we demonstrate the function of an Aβ-response synNotch receptor Bap-Notch in human iPSC-derived astrocytes, show proof-of-concept function in immortalized human microglia, and deliver Bap-Notch MSCs to aged 5xFAD mice. Additionally, we demonstrate that Bap-Notch astrocytes can express therapeutic transgenes that have effects independent of Aβ clearance – including an anti-inflammatory transgene cassette containing IL-1Ra and sTNFR1, which significantly attenuates reactive astrocyte gene expression.In conclusion, this work integrates a cell control module into a precision regenerative medicine strategy to treat AD. We have generated cells that can detect AD pathology and, in response, regulate therapeutic transgene expression. This platform functions both in brain-resident cells (here, microglia and astrocytes) and a clinical AD cell therapy candidate (MSCs), and thus represents a significant advance in the potential therapies for AD. The modularity of the system allows for flexibility in the choice of targeted transgene therapeutics. In all the AD drugs approved in the last 25 years, including Leqembi (lecanemab) and Kisunla (donanemab), the therapeutic benefit is reliant on clearance of Aβ protein aggregates. Such anti-Aβ monoclonal antibodies have demonstrated extremely limited cognitive benefit and are associated with potentially serious side effects, despite their ability to engage Aβ. Our approach builds upon the massive research effort to discover such anti-Aβ antibodies by repurposing them as the recognition domains of synthetic receptors. Crucially, we selected transgenes to address aspects of disease (BDNF, inflammatory cytokine antagonists) that intersect with facets of AD pathology that are not resolved by simply clearing Aβ from the AD brain (synapse and neuron loss; neuroinflammation; astrocyte-mediated BBB disruption). Further, the platform affords for autoregulation of therapeutic factors on the basis of local amyloid disease burden, thereby potentially avoiding complications from wide-spread, unregulated production of biologic drugs throughout the brain. Thus, this work represents a powerful advance over traditional therapies or investigative cell-based/gene therapies applied to AD, which lack a means for feedback-controlled cell functions based on pathology within the AD niche.

## Methods

### Receptor construction and plasmid cloning

Each Aβ receptor was constructed by reverse translating the amino acid sequences of the variable heavy (vH) and variable light (vL) chains of their respective monoclonal antibodies (donanemab, bapineuzumab, and gantenerumab). The scFv nucleic acid sequences were designed by combining cognate vH and vL chains via a flexible linker ((G_4_S)_3_). These scFvs were assembled with the Notch transmembrane core and tetracycline transactivator (tTA) synthetic transcription factor. In a series of experiments, a previously described, GFP-responsive receptor based on the GFP-specific LaG16 nanobody was used as a control.^67,96^ Lentiviral plasmids encoding for each receptor were cloned using NEBuilder HiFi DNA Assembly mix. Payload plasmids encoding the response transgene downstream of the tTA response element (TRE) were cloned using the same methods. For circuits where receptor activation resulted in expression of multiple transgenes (*e.g.* sTNFR1, IL-1Ra), transgenes were expressed from a biscistronic cassette using either a P2A ribosomal skipping peptide or an internal ribosomal entry site (IRES). Plasmids containing the LaG16 GFP-responsive synNotch receptor (Addgene 223536) and firefly luciferase-P2A-mCherry (Addgene 223539) and SEAP-IRES-mCherry payloads (Addgene 223535) have been previously used.^68^ Plasmids were transformed into DH5α *E. coli* competent cells (SMObio) and plated on LB agar with ampicillin and incubated overnight at 37°C. Colonies were picked and cultured in LB with ampicillin overnight. Plasmids were purified via miniprep (Qiagen). All plasmids were sequence-verified before use. Plasmids and their associated Addgene depository numbers are listed in Supplementary Table 1.

### Cell culture

#### L929 mouse fibroblasts

Fibroblasts (ATCC) were cultured in DMEM + GlutaMax (Gibco) supplemented with 10% heat-inactivated FBS (Gibco) at 37°C with 5% CO_2_. For routine passaging, cells were dissociated using TrypLE (Gibco) by incubation at 37°C for 5 minutes. For synNotch activation experiments, cells were detached using Accutase (Gibco) by incubation at 37°C for 5 minutes. Suspensions were quenched in medium and centrifuged at 300x*g* for 5 minutes and resuspended for plating.

#### Mouse mesenchymal stem cells

Bone marrow-derived MSCs from C57BL/6 mice (Cyagen) were cultured in MEM-α + GlutaMax (Gibco) supplemented with 15% FBS (Gibco) at 37°C with 5% CO_2_. For passaging, cells were washed with DPBS (Gibco) then detached with TrypLE by incubation at 37°C for 5 minutes. For transplanting to slice cultures, cells were detached using Accutase (Gibco) by incubation at 37°C for 5 minutes. Suspensions were quenched in medium and centrifuged at 300x*g* for 5 minutes and resuspended for seeding.

#### Human immortalized microglia

Human microglia immortalized via hTERT overexpression (ABM Goode) were cultured in Prigrow III medium (ABM Goode) with 10% heat-inactivated FBS (Gibco) at 37C with 5% CO2. For routine passaging, cells were washed with DPBS and disassociated using TrypLE by incubation at 37°C for 5 minutes. For synNotch activation experiments, cells were detached using Accutase by incubation at 37°C for 5 minutes. Suspensions were quenched in medium and centrifuged at 300x*g* for 5 minutes and resuspended for seeding.

#### CC3 human induced pluripotent stem cells (hiPSCs)

CC3 hiPSCs^97^ were maintained in Essential 8 (E8) medium^98^ on Matrigel (Corning)-coated wells. Routine cell passaging was carried out by incubation with ReLeSR (Stem Cell Technologies) for 1 minute at RT, followed by ReLeSR removal and incubation at 37°C for 6 minutes.

#### Astrocyte differentiation from CC3 hiPSCs

Differentiations were adapted from a previously described protocol.^21^ On Day −1 of differentiation, CC3 hiPSCs were detached using Accutase and plated on Matrigel in E8 with 10 µM Y-27632 (ROCKi) (Tocris) at a density of 2×10^5^ cells/cm^2^ (Fig. 3A). The following day, the medium was replaced with E6 containing 10 µM SB431542 (STEMCELL Technologies) and 1 µM dorsomorphin (Tocris). Medium was changed every day until Day 6 of differentiation. On Day 6, clumps of cells were carved out from the confluent cell layer using a P200 pipette tip and replated on a fresh Matrigel well containing E6 with 10 ng/mL CNTF (Peprotech) and 10 ng/mL EGF (Peprotech). The plates were shaken at 37°C for several hours to allow cells to attach. The medium was replaced every 3 days and cells were passaged when nearing confluence, with the first passage occurring around Day 30 of differentiation. To reach astrocyte maturity, cells were maintained in differentiation medium for 7 to 10 passages until about 65 days of culture. Astrocytic phenotype was confirmed by GFAP expression by immunocytochemistry.

### Lentivirus production and transduction

Lentivirus was produced by transfecting Lx293T cells (Clontech) plated in a 6-well plate with 2.0 µg of transfer plasmid (i.e., receptor or payload plasmids), 1.5 µg of pCMV-dR8.91 gag/pol packaging plasmid,^99^ and 0.6 µg of pMD2.G envelope plasmid (Addgene #12259) with Lipofectamine 3000 (Thermo Scientific). One day following transfection, the medium was replaced with complete medium composed of DMEM + GlutaMax supplemented with 10% heat-inactivated FBS (Gibco). On days 2 and 3 following transfection, viral medium was collected and filtered with a 0.45 µm PVDF filter (CELLTREAT).

#### Transduction of L929 fibroblasts and MSCs

Viral medium was concentrated in a 100 kDa MWCO filter (Millipore) via centrifugation and resuspended in fresh cell culture medium (DMEM + 10% heat-inactivated FBS; MEM-α + 15% FBS) for cell transduction. Cells were reverse-transduced in a 6-well plate format using the virus collected from one well. Media were supplemented with 4 µg/mL polybrene (Sigma Aldrich) to facilitate viral transduction. Media were changed the following day to remove virus.

#### Transduction of immortalized human microglia

Viral medium was incubated overnight at 4C with Lenti-X concentrator (Takara) and centrifuged at 1500xg for 45 minutes at 4C. Pelleted virus was resuspended in microglial medium. Cells were transduced in a 6-well plate format using the virus collected from one well. Media were supplemented with 4 µg/mL polybrene (Sigma Aldrich) to facilitate viral transduction. Cells were transduced using lentiviral spinoculation at 1000x*g* for 1 hour at room temperature followed by overnight incubation at 37°C. Media were changed the following day to remove virus.

#### Transduction of hiPSC-derived astrocytes

Viral medium was incubated overnight at 4°C with Lenti-X concentrator (Takara). Lentivirus was pelleted via centrifugation and resuspended in astrocyte medium (E6 + CNTF + EGF) before being added to astrocytes. Cells were transduced in a 6-well plate format using the half the virus collected from one well. Media was supplemented with 4 µg/mL polybrene (Sigma Aldrich) to facilitate viral transduction. Media were changed the following day to remove virus.

### Flow cytometry and fluorescence activated cell sorting (FACS)

Prior to analytical flow cytometry or FACS, cells were dissociated with Accutase. Cells were centrifuged at 300x*g* for 5 minutes and resuspended in flow buffer (1% BSA in DPBS) and blocked on ice for 15 minutes. Cells were stained with an anti-c-myc tag antibody conjugated to Alexa647 (Cell Signaling Technologies) diluted 1:50 in flow buffer for 1 hr on ice. Cells were washed 2 times by centrifugation and resuspended in flow buffer for analysis on a Cellstream flow cytometer to assess cells double-positive for myc-tag (incorporated on the N-terminus of the synNotch receptor) and BFP (encoded for constitutive expression in the payload transgene vector). Flow cytometry analysis was done using FlowJo. To obtain a pure population of synNotch-positive cells, samples were run on a 4-laser FACSAria III cell sorter, and double-positive cells were collected. Collected cells were spun down at 300x*g* for 5 minutes and resuspended in cell culture medium for plating.

### Synthetic Aβ preparation

Biotinylated and unmodified synthetic Aβ_40_ and Aβ_42_ preparations (Anaspec) were resuspended according to manufacturer’s instructions and aliquoted for long-term storage at −80°C. Protein was originally resuspended at 1 mg/mL for dynamic light scattering (DLS) analysis then diluted to 50 µg/mL for cell culture assays.

SynNotch L929 activation

For immobilized, biotinylated Aβ, tissue culture plates were treated with 10 µg/mL of streptavidin (Thermo Fisher) in DPBS and incubated for a minimum of 1 hr at 37°C. The streptavidin was then removed, and biotinylated Aβ was added to the coated wells at 50 µg/mL and incubated for an additional hour at 37°C. The Aβ solution was removed and rinsed with DPBS, leaving 0.6 µg Aβ per well by BCA. SynNotch L929 cells were then plated at 20,000 cells/well. For adsorbed unmodified Aβ, the cell culture plate was coated directly with 50 µg/mL Aβ and incubated for 1 hr at 37°C. The wells were rinsed and cells were added as above. For medium-supplemented Aβ, the cells were first plated at 20,000 cells/well and 50 µg/mL of Aβ was supplemented to the medium. After 72 hr, the production of the firefly luciferase transgene was measured using a BrightGlo luminescence assay (Promega) on a Tecan Infinite M1000 Pro plate reader.

### Immunostaining of astrocytes

Immunolabeling of hiPSC-derived astrocytes was performed at day 60 of differentiation to confirm fate. Wells were fixed with ice-cold 4% paraformaldehyde in DPBS (Thomas Scientific) for 15 minutes at room temperature. Cells were then washed 3 times with DPBS and blocked with 5% FBS, 0.3% Triton-X in DPBS at 4°C overnight. Cells were incubated with anti-GFAP (Aves Labs, 1:300) and anti-CD44 (Cell Signaling Technologies, 1:500) primary antibodies at 4°C overnight. After primary antibody incubation, cells were rinsed 3 times with DPBS and then incubated with secondary antibodies for 1 hr at room temperature (goat anti-chicken Alexa488; goat anti-mouse Alexa647, Invitrogen, 1:1000). Cells were then incubated with nuclear stain DAPI (Thermo Scientific) and washed before imaging.

### Human brain-derived Aβ seed preparation

Brain tissue from de-identified human cases with cerebral amyloid angiopathy (CAA) pathology but minimal parenchymal Aβ and tau (i.e., CAA-Aβ) from the Vanderbilt Brain and Biospecimen Bank (VBBB) were used to isolate Aβ seeds. Ethical oversight of the VBBB is provided by the Vanderbilt Institutional Review Board; recruitment and uses of the tissue conforms to the ethical principles of the Belmont Report. Microvessels were isolated from identified brains and Aβ microparticles ranging from 500 nm to 7 µm in diameter were collected via collagenase digestion.^100^ Aβ was then separated from the vascular elements using a 3D printed custom-designed micro-sieve. Western blots were performed on samples to confirm that they were not contaminated by tau.

### SynNotch astrocyte activation on CAA-Aβ

SynNotch astrocytes were plated on Matrigel-coated wells at a density of 50,000 cells/cm^2^. 0.6 µg of CAA-Aβ seeds was added directly to plated astrocytes. To prevent contamination by the patient-derived protein, cultures were maintained in 1X antibiotic-antimycotic (Gibco). SynNotch activation was visualized by mCherry by fluorescence microscopy after 72 hr. Medium was collected for detection of secreted embryonic alkaline phosphatase (SEAP) by Great EscAPe SEAP Chemiluminescence Kit (Takara) using a Tecan Infinite M1000 Pro plate reader. Astrocyte number was normalized by adding 1X CellTiter-Fluor reagent (Promega) to wells after medium collection; fluorescence was measured at Ex 390 nm/Em 505 nm on the same plate reader.

### SynNotch-BDNF astrocyte activation on synthetic Aβ

Stem cell-derived astrocytes are typically cultured on a basement membrane such as Matrigel. For the BDNF-expressing astrocytes, cells were cultured on a surface decorated with glycosaminoglycan-binding peptide (GBP) and cyclic arginine-asparagine-aspartic acid (cRGD) as a as a substitute for Matrigel to promote cell adhesion.^96,101^ Thus, biotinylated GBP (Genscript Express) and cRGD (Carbosynth) adhesion peptides were reconstituted in UltraPure distilled water (Invitrogen) and mixed with biotinylated Aβ_42_ to final concentrations of 5 µM GBP, 2.15 µM cRGD, and 0.4 µM of Aβ. Non-tissue culture treated well plates were first coated with 10 µg/mL of streptavidin for a minimum of 1 hr to overnight at 37°C. The streptavidin was aspirated and replaced with the peptide solution and incubated for 1-2 hr at 37°C. SynNotch astrocytes were dissociated with Accutase and plated in the coated wells after the peptide solution was removed.

### Astrocyte monoculture inflammation experiments

Synthetic Aβ_42_ was incorporated into Matrigel working solution at a concentration of 50 µg/mL. Wells were coated with the Aβ-embedded Matrigel and incubated for 1 hr at 37°C. Control wells were coated with Matrigel without added Aβ. Matrigel solution was removed and cells were plated in astrocyte medium. SynNotch astrocytes were dissociated with Accutase and plated at a density of ∼35,000 cells/cm^2^. 48 hr after cell plating, the cells were supplemented with inflammatory cytokines IL-1α (5 ng/mL, STEMCELL Technologies) and TNF-α (10 ng/mL, STEMCELL Technologies). After an additional 48 hr, media were collected for analysis by ELISA or SEAP assay and mRNA was isolated. The presence of the SEAP transgene was detected using a SEAP activity assay ((Great EscAPe Chemiluminescence Kit 2.0, Takara) measured on a Tecan Infinite M1000 Pro plate reader.

### ELISA

The Human TNF R1/TNFRSF1A DuoSet ELISA kit and Human Il-1RA/IL-1F3 DuoSet ELISA kits were used with the DuoSet Ancillary Reagents kit to detect anti-inflammatory transgenes (R&D). Cell culture media from Bap-Notch_sTNFR1-IL-1Ra astrocytes were diluted 1:50 in diluent for sTNFR1 ELISA, 1:100 for IL-1Ra ELISA; media from Bap-Notch_SEAP astrocytes were diluted 1:5 in diluent. For detection of BDNF payload, the Total BDNF Quantikine ELISA kit was used (R&D). Cell culture media were added directly with no dilution. Absorbance was measured at 450 nm on a Tecan Infinite M1000 Pro plate reader and a correction read at 570 nm was subtracted. The average absorbance of the 0 pg/mL standard was subtracted from all samples. Sample concentrations were calculated using a four-parameter logistic curve fit of the standards using Graphpad Prism 10.

### Gene expression analysis

Cells were lysed, and mRNA was isolated using PureLink Mini Kits (Invitrogen) following the manufacturer’s instructions. mRNA was reverse transcribed to cDNA using the SuperScript IV VILO Master Mix (Invitrogen). Quantitative PCR was performed with PowerTrack SYBR Green Master Mix (Applied Biosystems) on a Bio-Rad CFX96 with the mass of cDNA held constant across all samples. Primer pairs used for gene expression detection are listed in Supplementary Table 2. Relative gene expression was calculated using the delta-delta Ct method using GAPDH as a reference gene. In all inflammation experiments, samples from the Aβ-free/cytokine-free condition were used for the reference group.

### 5xFAD animal husbandry

All animal protocols were approved by the Institutional Animal Care and Use Committee (IACUC) at Vanderbilt University. Mice were housed and bred as described previously.^102^ Briefly, male hemizygous 5xFAD mice (The Jackson Laboratory RRID: MMRRC_03848-JAX) and female wild-type C57BL/6J (RRID: IMSR_JAX:000664) were bred to establish paternal inheritance. Mice were genotyped by tail biopsy using Transnetyx probes for huPSEN, APPsw, Chr3-6_WT).

### Organotypic slice culture preparation and cell transplantation

All animal protocols were approved by the Institutional Animal Care and Use Committee (IACUC) at Vanderbilt University. Three adult (11-month old) hemizygous 5xFAD mice and 3 adult (11-month old) wild-type C57/BL6 littermates mice were sacrificed. Brains were removed and mounted on a vibrating blade microtome (Leica) in slicing media composed of Hibernate A (BrainBits) with B27, GlutaMax, and Gentamycin (Gibco). Coronal slices 300 µm thick were cultured on a membrane insert (Millipore Sigma) in Neurobasal A (Gibco) with B27, GlutaMax, and Gentamycin. After slices were prepared, synNotch MSCs were dissociated with Accutase and resuspended in MEM-α with 15% FBS. 200,000 synNotch cells were plated in a drop of 10 µL on top of each slice. Save for a few slices maintained as no-MSC controls, half the slices from each mouse received Bap-Notch MSCs; half received control LaG16-Notch MSCs. Serial slices were alternately seeded with Bap-Notch MSCs or LaG16-Notch MSCs to remove the confounding effects of variable Aβ accumulation on different sections of the brain. After 72 hr, medium was collected from the culture and the presence of SEAP transgene was determined using a chemiluminescence SEAP activity assay (Great EscAPe Chemiluminescence Kit 2.0, Takara) read on a Tecan Infinite Pro plate reader. Samples were run in technical duplicate.

### Organotypic slice culture cryosection preparation and staining

Slices were fixed in 4% paraformaldehyde at 4°C overnight and cryopreserved in 30% sucrose. Samples were embedded in optimal cutting temperature (OCT) compound and tissue was sectioned into 10 µm sections. After thaw, sections were surrounded with a hydrophobic barrier, washed with DPBS, and blocked with 1% BSA, 0.25% Triton-X, and 10% normal goat serum for 1 hr at RT. Sections were incubated with anti-Aβ (6E10, 1:400) and anti-mCherry (Cell Signaling Technologies, 1:200) primary antibodies at 4°C overnight. After primary antibody incubation, cells were rinsed 3 times with DPBS and incubated with secondary antibodies for 1 hr at room temperature (goat anti-mouse Alexa488 (1:1000); goat anti-rabbit Alexa555 (1:500) (Invitrogen). After secondary antibody incubation, tissues were incubated with TrueVIEW Autofluorescence Quenching reagent (VectorLabs) for 5 min at RT. Finally, sections were washed and sealed with a coverslip. Sections were cured for 2 hours at RT prior to imaging.

### SynNotch MSC transplantation to live animals

Three adult (12-month old) hemizygous 5xFAD mice and 3 adult (12-month old) wild-type C57/BL6 mice were injected with synNotch cells. Mice were anesthetized by 2-3% isoflurane inhalation and placed in a stereotactic rig. Each mouse was injected bilaterally with 150,000 Bap-Notch MSCs in 2 µL sterile PBS per injection. Prior to injection, Bap-Notch MSCs were sorted for double positive GFP (payload vector) and Myc-Tag on the receptor). Cells were injected using coordinates (−2.06 AP, +/-1.75 ML, −1.75 DV relative to bregma).^103^ Cells were injected at a rate of 0.35 µL/second. After the entire 2 µL volume was dispensed, the needle was left in place for an additional 5 minutes before withdrawal. After surgery, mice were treated with 5 mg/kg of Ketoprofen every 24 hr. After 72 hr, mice were anesthetized via intraperitoneal Avertin injection (250 mg/kg). CSF draws were performed by making an incision at the back of the neck to expose the cisterna magna and extracting CSF using a catheter consisting of a 30-gauge needle attached to a 25 µL Hamilton syringe slowly over 3-5 minutes. CSF was visually confirmed not to be contaminated by blood. After CSF collection, blood was drawn by cardiac puncture, allowed to clot, and centrifuged at 10,000xg for 10 min to allow for serum collection off the top. CSF and serum samples were stored at −80°C until analysis. Mice were finally perfused with PBS followed by 10% formalin. Whole brains were fixed in 10% formalin for 24 hr, followed by sucrose gradient (10%, 20%, 30% until tissue sinking) cryopreservation. Brains were embedded in OCT and cryosectioned at 10 µm. Cryosections were stained using the same process as above for Aβ (6E10, 1:400 with Invitrogen goat-anti-mouse Alexa 405, 1:1000), mCherry (CST, 1:200; goat-anti-rabbit Alexa647 1:1000), and GFP (ChromoTek GFP-Booster Alexa 488). CSF and serum was analyzed using chemiluminescence assay read on a Tecan Infinite Pro plate reader. Serum samples were run in technical duplicate and serum from uninjected aged (∼12-months-old) 5xFAD (2 female; 1 male) and WT (3 male) mice were included as controls. Due to volume considerations, CSF samples were run in singlicate using 5 µL/sample (all other SEAP assays followed kit recommended volume of 25 µL/sample).

### Fluorescence microscopy

All images were taken on a Leica Dmi8 epifluorescent microscope at 10 or 20X magnification.

### Statistical analysis

All statistical analyses were performed with GraphPad Prism 10. Plotted values represent mean ± standard deviation. Outliers were identified by robust regression and outlier removal (ROUT) and were excluded. Statistical analysis of qPCR gene expression was performed on the log_2_ transform of fold change. Non-parametric statistical analyses were used for the *in vivo* data in Fig. 6. Parametric tests were used for all other analyses, and non-normal data were logtransformed before statistical analysis. For all analyses, α=0.05. All comparative statistics, including t and F statistics, degrees of freedom, P values, and effect size are included in the supplementary information.

## Acknowledgements

FACS was performed at the Vanderbilt Flow Cytometry Shared Resource, which is supported by the Vanderbilt Ingram Cancer Center (P30 CA068485) and the Vanderbilt Digestive Disease Research Center (DK058404). We thank Zach Lamantia for technical assistance.

## Author contributions

MRS, ESL, and JMB conceived of the study and designed all experiments. MRS, SRS, HJB, DJC, and RSB carried out all experiments. HK provided iPSC-derived astrocytes and technical assistance with subsequent astrocyte differentiations. APL performed all stereotactic injections and CSF collections. DC provided key input for astrocyte experiments. HJB assisted with synthetic Aβ preparation and delivery to astrocytes. RJB performed all 5xFAD breeding and husbandry. AS and MSS provided human tissue samples. MRS and JMB wrote the manuscript.

