## Supplementary Material for "Amyloid-β-regulated gene circuits for programmable Alzheimer’s disease therapy"

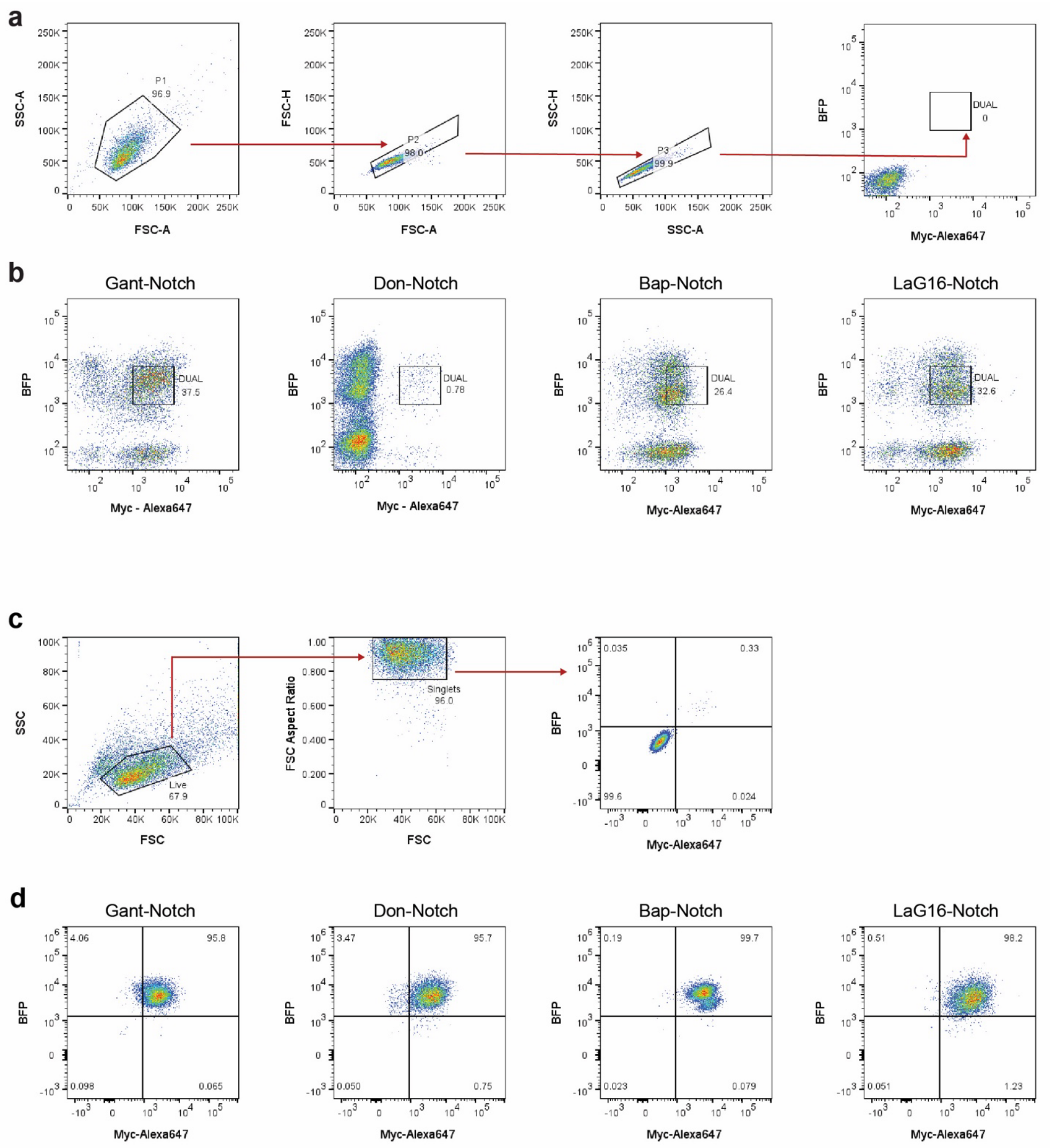


**Supplementary Figure 1.** FACs plots for synNotch L929s. (A) Gating strategy for live cells, singlets, and synNotch positive cells on non-transduced L929s. (B) Population of cells sorted for Gant-Notch, Don-Notch, and Bap-Notch L929 cell lines. Cells double-positive for BFP (payload vector) and the myc-tag (receptor construct) were sorted at the same level for the full panel of Aβ-synNotch fibroblasts. (C) Analytical flow gating strategy for live cells, singlets and synNotch positive cells on non-transduced L929s. (D)

SynNotch-positive L929 cells 8 months after cell sorting. Cells double-positive for BFP (payload vector) and the myc-tag (receptor construct) are considered synNotch-positive.

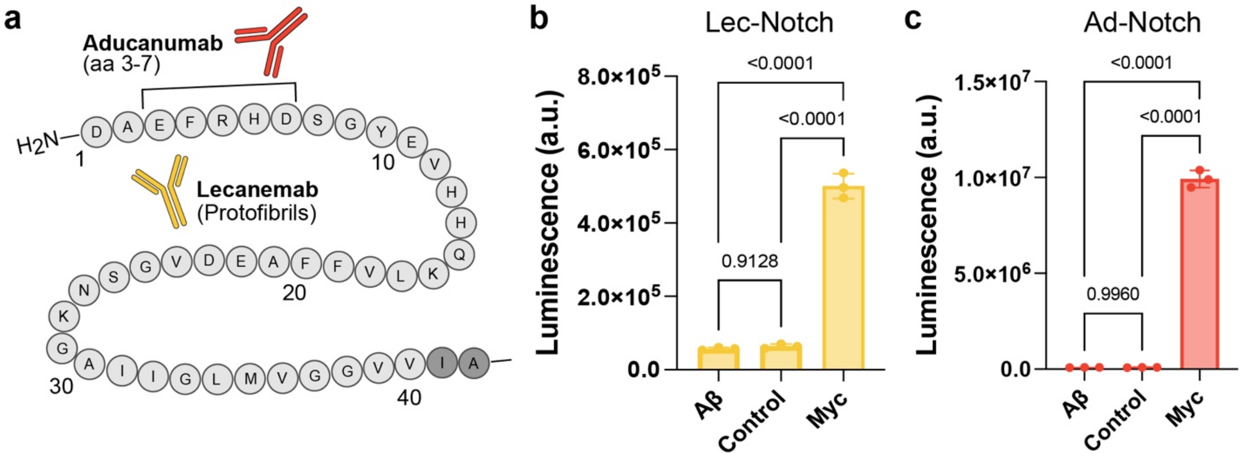


**Supplementary Figure 2.** Testing of Ad-Notch and Lec-Notch receptors. (A) Depiction of aducanumab and lecanemab epitopes on the Aβ peptide. Aβ-driven luciferase transgene expression of (B) Lec-Notch or (C) Ad-Notch L929 cells plated on biotinylated $A\beta_{42}$ or cultured with 1 µm beads decorated with antibodies specific to the c-myc epitope tag. $n$ = 3; mean±SD; stats from one-way ANOVA with Tukey's multiple comparisons.

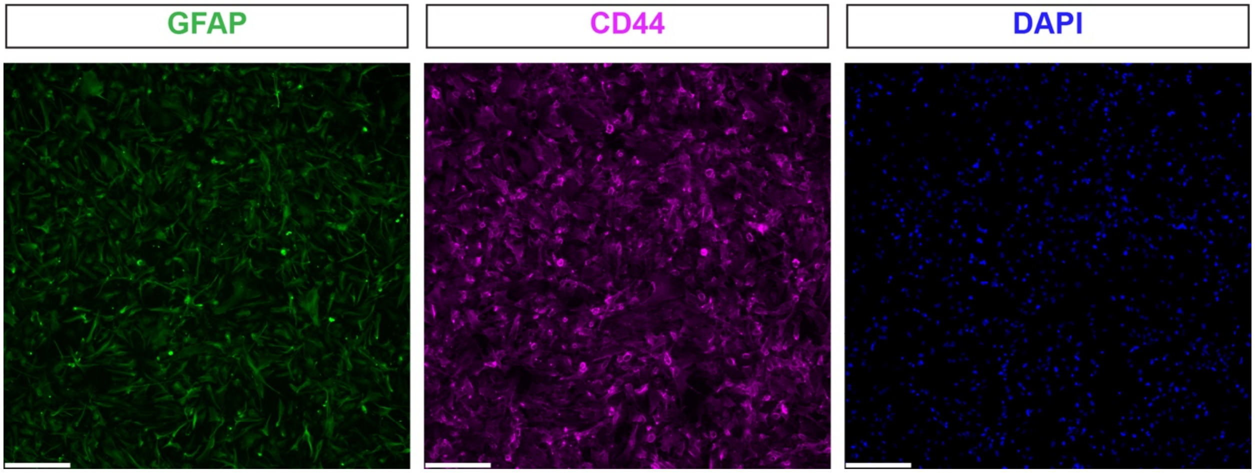


**Supplementary Figure 3.** Astrocyte marker expression in differentiated cells. GFAP (green) and CD44 (magenta) expression by ICC. DAPI shows cell nuclei. Scale bar = 200 µm. Merged image shown in Fig. 3D.

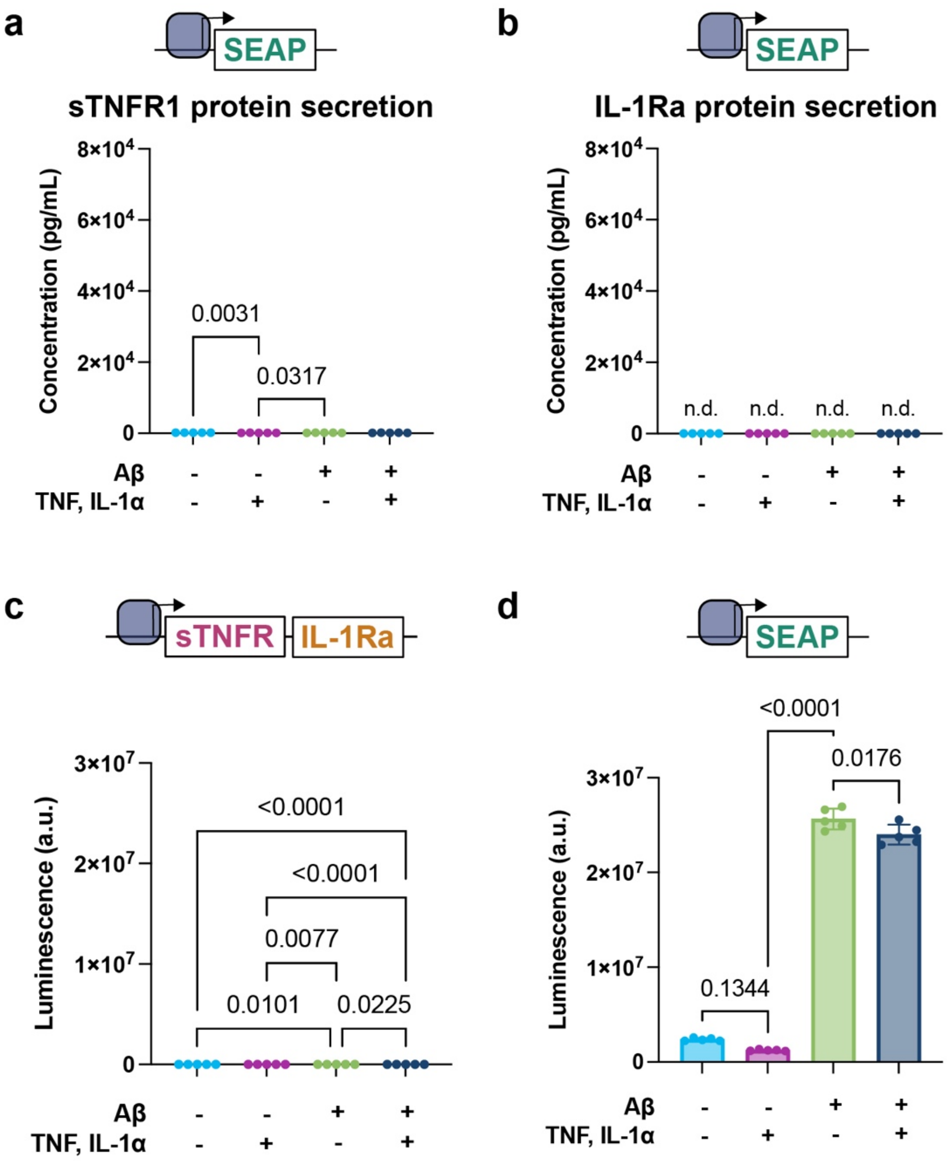


**Supplementary Figure 4.** Regulation of Aβ-driven transgene expression in Bap-Notch astrocytes. sTNFR1 (A) and IL-Ra (B) ELISA on medium of Bap-Notch astrocytes engineered to express SEAP in response to Aβ. Astrocytes are plated on immobilized $A\beta_{42}$ and treated with TNF and IL-1α. *n* = 5; mean±SD; stats from two-way ANOVA with Tukey's multiple comparisons. n.d. = not detected. (C) Aβ-driven SEAP expression of Bap-Notch astrocytes programmed with either sTNFR1-IL-1Ra or SEAP payloads. *n* = 5; mean±SD; two-way ANOVA with Tukey's multiple comparisons.

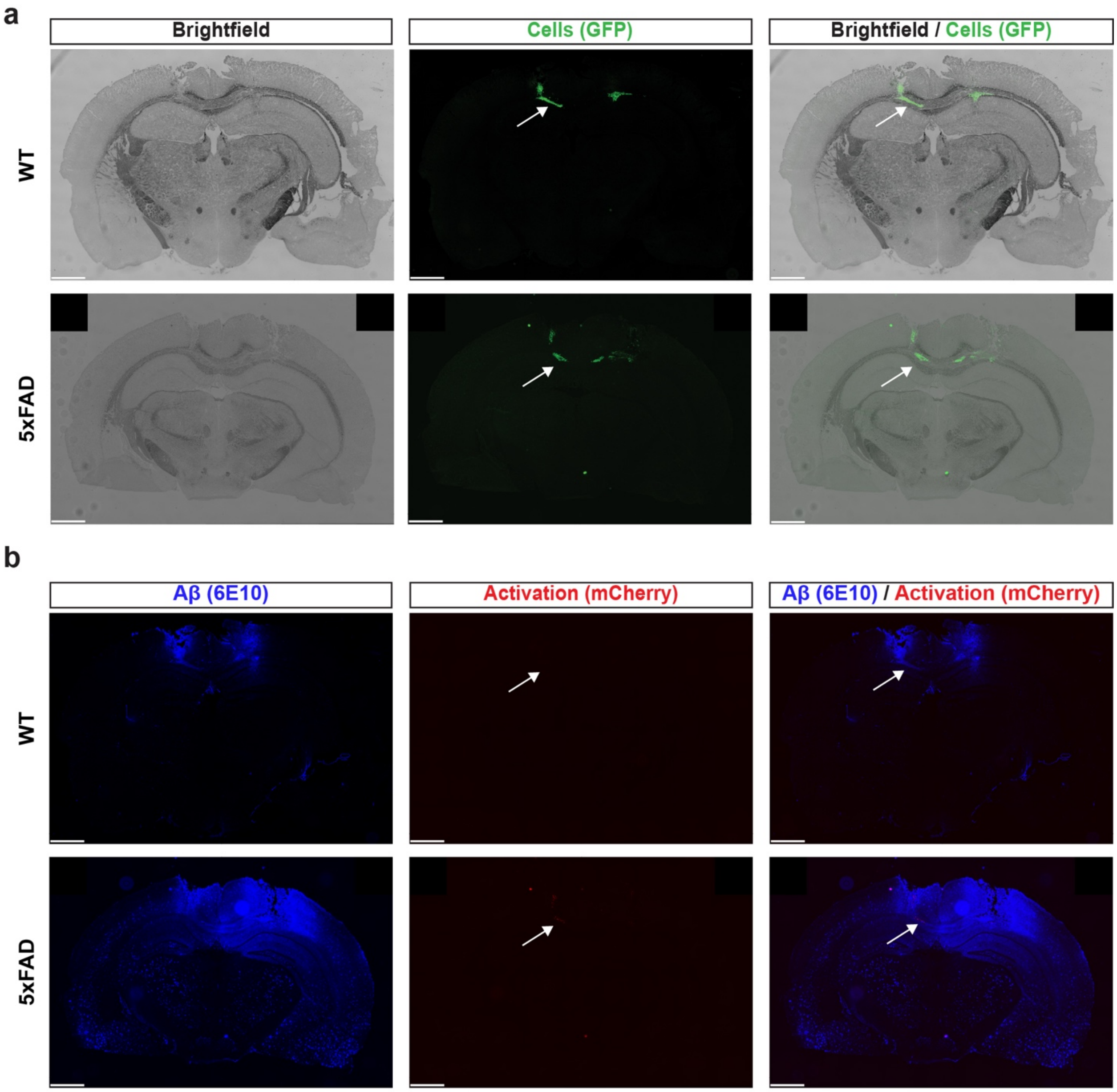


**Supplementary Figure 5.** Whole brain scans of mice injected with Bap-Notch MSCs. (A) Brightfield and fluorescence microscopy showing location of injected Bap-Notch MSCs (GFP+) in WT and 5xFAD brains. Scale bar = 1 mm. (B) Fluorescence microscopy showing Aβ deposition (6E10) and Aβ-driven synNotch activation (mCherry) in 5xFAD brain. Scale bar = 1 mm. Arrows indicate regions shown at higher magnification in main Figure 6.

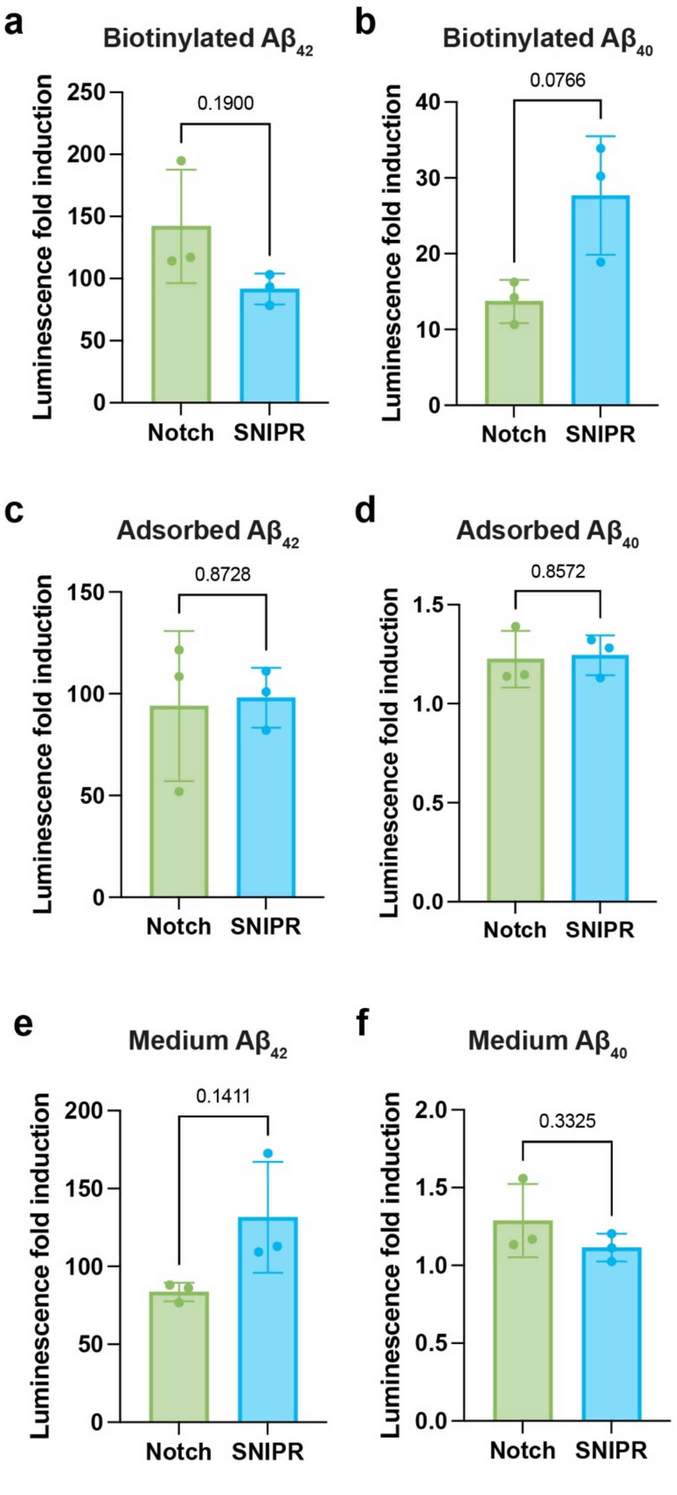


**Supplementary Figure 6.** Bap-SNIPR and Bap-Notch comparisons. Aβ-driven luciferase transgene expression of L929 fibroblasts expressing Bap-Notch and Bap-SNIPR plated on biotinylated $A\beta_{42}$ (A) or $A\beta_{40}$ (B), adsorbed $A\beta_{42}$ (C) or $A\beta_{40}$ (D), or medium-supplemented $A\beta_{42}$ (E) or $A\beta_{40}$ (F) $n$ = 3; mean±SD; Welch's $t$-test.

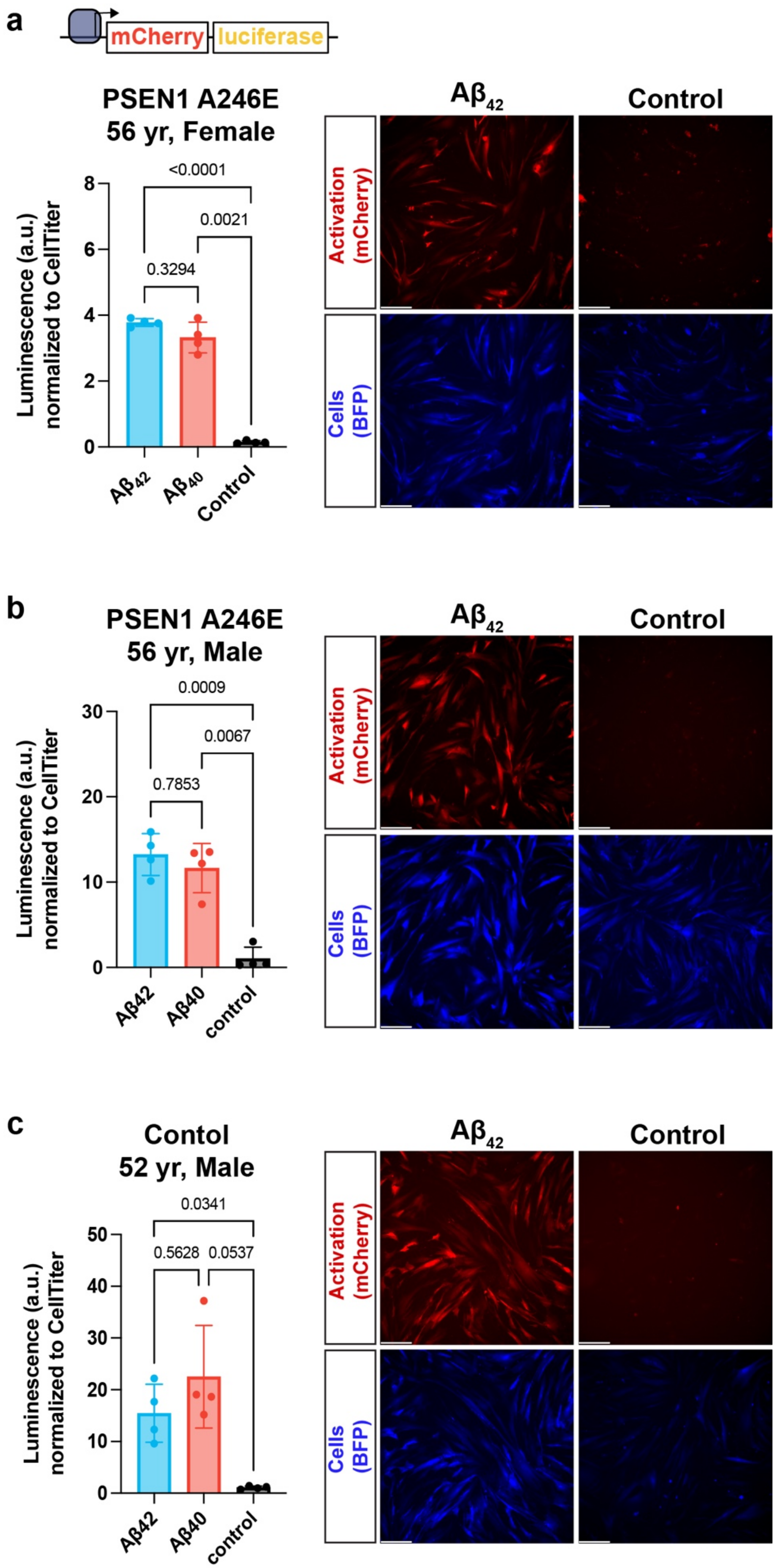
a
mCherry
luciferase
PSEN1 A246E
56 yr, Female
Luminescence (a.u.) normalized to CellTiter
<0.0001
0.0021
0.3294
Aβ42
Aβ40
Control
Activation (mCherry)
Cells (BFP)
b
PSEN1 A246E
56 yr, Male
0.0009
0.0067
0.7853
Aβ42
Aβ40
control
c
Contol
52 yr, Male
0.0341
0.5628
0.0537

**Supplementary Figure 7.** Testing of Bap-Notch in PSEN1 patient-derived fibroblasts. Aβ-driven luciferase and mCherry transgene expression of Bap-Notch fibroblasts collected from a 56-year-old female with a missense PSEN1 mutations and early-onset AD (A), a 56-year-old male with a missense PSEN1 mutation (B), and a 52-year-old apparently healthy male (C). $n$=4; mean mean±SD; stats from Welch ANOVA with Dunnett's T3 multiple comparisons. BFP is a constitutive cell marker. Scale bar = 200 μm.

### Supplementary Methods

Anti-c-myc bead activation

Lec-Notch and Ad-Notch L929s were activated in an Aβ-independent manner by adding anti-c-myc beads at a final concentration of 0.1 mg/mL (Thermo Fisher). Aβ recognition was tested using biotinylated $A\beta_{42}$ with the same methods used for Gant-Notch, Don-Notch, and Bap-Notch (Fig. 1; Main methods; SynNotch L929 activation). Luciferase activity was measured after 72 hr with a luminescence assay (Promega) as noted previously.

Aβ western blot

Synthetic Aβ was prepared as described in the main text, frozen at -80˚C and was thawed directly for assessment by western blot and DLS. 100 ng of protein with native sample buffer was loaded in wells of a 4-20% PAGE gel (Biorad), run in Tris-Glycine buffer without SDS, and transferred onto a nitrocellulose membrane. The blot was blocked using SuperBlock T20 Blocking Buffer (Thermo Fisher) and then incubated with the 6E10 anti-Aβ antibody (Biolegend, 1:1000 dilution). The next day, the membrane was washed 3 times rapidly, followed by 3x 5-minute washes with Tris Buffered Saline with Tween 20 (TBST). The membrane was incubated with goat anti-mouse IgG HRP (BioRad, 1:5,000) for 1 hr at room temperature and washed. The western blot was visualized using an Odyssey Fc (LI-COR).

Dynamic light scattering (DLS)

Size measurements of synthetic Aβ aggregates were performed on a Malvern Zetasizer in the Vanderbilt Institute for Nanoscale Science and Engineering (VINSE) core facility using a low volume disposable sizing cell and standard parameters. An attenuation between 7 and 10 was selected for all Aβ samples. Aβ samples were run at concentrations of 1 mg/mL, 0.5 mg/mL, and 0.25 mg/mL.

Patient-derived fibroblast

Fibroblasts collected via skin biopsy from subjects with cognitive decline due to a missense *PSEN1* mutation (A246E) (56-year-old man, AG06840; 56-year-old woman AG06848) and from an apparently healthy subject (52-year-old man GM23967) were purchased from Coriell Institute and were cultured according to instructions. Cells were maintained in Eagle's Minimum Essential Medium with Earle's salts, 15% FBS, non-essential amino acids, and 2 mM L-glutamine. Upon reaching confluency, cells were passaged 1:2. For transduction, viral medium was incubated overnight at 4˚C with Lenti-X concentrator (Takara). Lentivirus was pelleted via centrifugation and resuspended in fibroblast medium. Cells were plated in a 6-well plate at a density of ~15k/cm$^2$ and were transduced the following day by adding virus with 4 μg/mL polybrene. Media were changed the following day to remove virus. Transduced cells were selected via puromycin at 4 μg/mL before plating the cells for Aβ detection. For Aβ detection, tissue culture plates were treated with 10 μg/mL of streptavidin (Thermo Fisher) in DPBS and incubated for a minimum of 1 hr at 37˚C. The streptavidin was then removed, and biotinylated Aβ was added to the coated wells at 50 μg/mL and incubated for an additional hour at 37˚C. The Aβ solution was removed and rinsed with DPBS and fibroblasts were plated at a density of ~46k/cm$^2$. SynNotch activation was visualized by mCherry by fluorescence microscopy. After 72 hr, the production of the firefly luciferase transgene was measured using a BrightGlo luminescence assay (Promega) on a Tecan Infinite M1000 Pro plate reader. Cell number was normalized by adding 1X CellTiter-Fluor reagent (Promega) to wells after medium collection; fluorescence was measured at Ex 390 nm/Em 505 nm on the same plate reader.

**Supplementary Table 1:** Description and Addgene Numbers for all plasmids used.

| Plasmid | Addgene Number |
|---|---|
| pLV EF1A_Gant-Notch-tTA_PGK_puroR | 261703 |
| pLV EF1A_Don-Notch-tTA_PGK_puroR | 261704 |
| pLV EF1A_Bap-Notch-tTA_PGK_puroR | 261705 |
| pLV EF1A_LaG16-Notch-tTA_PGK_puroR | 223536[1] |
| pLV TRE_FLuc-P2A-mCherry_PGK_BFP | 223539[1] |
| pLV gfaABC$_1$D_Bap-Notch-tTA_PGK_puroR | 261706 |
| pLV TRE_BDNF-P2A-mCherry_PGK_BFP | 261707 |
| pLV gfaABC$_1$D_SEAP | 261708 |
| pLV gfaABC$_1$D_sTNFR1-P2A-IL-1Ra | 261709 |
| pLV TRE_SEAP-IRES-mCherry_PGK_BFP | 223535[1] |
| pLV TRE_SEAP-IRES-mCherry_PGK_GFP | 261710 |
| pLV EF1A_Ad-Notch-tTA_PGK_puroR | 261711 |
| pLV EF1A_Lec-Notch-tTA_PGK_puroR | 261712 |
| pLV EF1A_Bap-SNIPR-tTA_PGK_puroR | 261713 |

**Supplementary Table 2:** Sequences of oligonucleotides used as primers for RT-qPCR gene expression analysis.

| Target | Sequence |
|---|---|
| *IL6* | For: 5’-AGACAGCCACTCACCTCTTCAG-3’<br>Rev: 5’-TTCTGCCAGTGCCTCTTTGCTG-3’ |
| *CSF2* | For: 5’-TCCTGAACCTGAGTAGAGACAC-3’<br>Rev: 5’-TGCTGCTTGTAGTGGCTGG-3’ |
| *C3* | For: 5’-GTGGAAATCCGAGCCGTTCTCT-3’<br>Rev: 5’-GATGGTTACGGTCTGCTGGTGA-3’ |
| *GAPDH* | For: 5’-GTCTCCTCTGACTTCAACAGCG-3’<br>Rev: 5’-ACCACCCTGTTGCTGTAGCCAA-3’ |

**Supplementary Table 3:** Comparative statistics for Figure 1. Grayed-out rows represent ANOVAs, with subsequent rows denoting multiple comparisons. ‡ represents data that were log transformed before statistical analysis.

| | Test/comparison | Statistic | df | P value | Effect size | 95% CI |
|---|---|---|---|---|---|---|
| Figure 1c | *Welch ANOVA with Dunnett's T3 multiple comparisons* | *W = 71.06* | *3.000, 20.01* | *<0.0001* | | |
| | Gant vs. Don | t = 3.588 | 20.34 | 0.0107 | mean difference = 28.47 | 5.477 to 51.46 |
| | Gant vs. Bap | t = 1.488 | 23.87 | 0.5965 | mean difference = -13.6 | -39.66 to 12.46 |
| | Gant vs. LaG16 | t = 8.211 | 12 | <0.0001 | mean difference = 54.98 | 34.26 to 75.70 |
| | Don vs. Bap | t = 5.577 | 21.21 | <0.0001 | mean difference = -42.07 | -63.82 to -20.31 |
| | Don vs. LaG16 | t = 6.226 | 12.01 | 0.0003 | mean difference = 26.51 | 13.33 to 39.69 |
| | Bap vs. LaG16 | t = 11.01 | 12 | <0.0001 | mean difference = 68.58 | 49.31 to 87.85 |
| Figure 1d | *Welch ANOVA with Dunnett's T3 multiple comparisons ‡* | *W = 200.2* | *3.000, 19.09* | *<0.0001* | | |
| | Gant vs. Don | t = 4.366 | 16.81 | 0.0025 | mean difference = 0.7497 | 0.2435 to 1.256 |
| | Gant vs. Bap | t = 1.262 | 22.09 | 0.7489 | mean difference = -0.1397 | -0.4575 to 0.1781 |
| | Gant vs. LaG16 | t = 14.03 | 12.22 | <0.0001 | mean difference = 1.255 | 0.9781 to 1.531 |
| | Don vs. Bap | t = 5.528 | 13.94 | 0.0004 | mean difference = -0.8894 | -1.376 to -0.4032 |
| | Don vs. LaG16 | t = 3.434 | 10.07 | 0.0342 | mean difference = 0.505 | 0.03445 to 0.9756 |
| | Bap vs. LaG16 | t = 21.04 | 12.4 | <0.0001 | mean difference = 1.394 | 1.189 to 1.600 |

**Supplementary Table 4:** Comparative statistics for Figure 2. Grayed-out rows represent ANOVAs, with subsequent rows denoting multiple comparisons.

| | Test/comparison | Statistic | df | P value | Effect size | 95% CI |
|---|---|---|---|---|---|---|
| Figure 2a | *Welch ANOVA with Dunnett's T3 multiple comparisons* | *W = 751.1* | *3.000, 17.52* | *<0.0001* | | |
| | Gant vs. Don | t = 11.61 | 19.71 | <0.0001 | mean difference = 69.92 | 52.47 to 87.37 |
| | Gant vs. Bap | t = 3.403 | 18.28 | 0.0182 | mean difference = -19.67 | -36.60 to -2.739 |
| | Gant vs. LaG16 | t = 27.22 | 9.001 | <0.0001 | mean difference = 122.1 | 107.5 to 136.8 |
| | Don vs. Bap | t = 16.52 | 22.93 | <0.0001 | mean difference = -89.59 | -105.1 to -74.08 |
| | Don vs. LaG16 | t = 13 | 12 | <0.0001 | mean difference = 52.22 | 39.79 to 64.65 |
| | Bap vs. LaG16 | t = 38.92 | 11 | <0.0001 | mean difference = 141.8 | 130.4 to 153.3 |
| Figure 2b | *Welch ANOVA with Dunnett's T3 multiple comparisons* | *W = 147.8* | *3.000, 20.00* | *<0.0001* | | |
| | Gant vs. Don | t = 1.977 | 21.05 | 0.2977 | mean difference = 17.49 | -8.031 to 43.02 |
| | Gant vs. Bap | t = 7.612 | 23.81 | <0.0001 | mean difference = -75.65 | -104.0 to -47.32 |
| | Gant vs. LaG16 | t = 7.239 | 12 | <0.0001 | mean difference = 53.11 | 30.40 to 75.81 |
| | Don vs. Bap | t = 11.18 | 22.09 | <0.0001 | mean difference = -93.14 | -117.1 to -69.21 |
| | Don vs. LaG16 | t = 7.193 | 12 | <0.0001 | mean difference = 35.61 | 20.29 to 50.93 |
| | Bap vs. LaG16 | t = 19.21 | 12 | <0.0001 | mean difference = 128.8 | 108.0 to 149.5 |
| Figure 2c | *Welch ANOVA with Dunnett's T3 multiple comparisons* | *W = 18.97* | *3.000, 20.07* | *<0.0001* | | |
| | Gant vs. Don | t = 4.102 | 13.54 | 0.0062 | mean difference = 5.488 | 1.444 to 9.532 |
| | Gant vs. Bap | t = 1.54 | 19.8 | 0.563 | mean difference = 2.336 | -2.061 to 6.733 |
| | Gant vs. LaG16 | t = 4.98 | 12.01 | 0.0018 | mean difference = 6.459 | 2.446 to 10.47 |
| | Don vs. Bap | t = 3.69 | 16.07 | 0.0114 | mean difference = -3.152 | -5.689 to -0.6155 |
| | Don vs. LaG16 | t = 2.941 | 12.1 | 0.0657 | mean difference = 0.971 | -0.05076 to 1.993 |
| | Bap vs. LaG16 | t = 5.23 | 12.02 | 0.0012 | mean difference = 4.123 | 1.683 to 6.563 |

| | | | | | | |
|---|---|---|---|---|---|---|
| Figure 2d | *Welch ANOVA with Dunnett's T3 multiple comparisons* | *W = 84.05* | *3.000, 23.53* | *<0.0001* | | |
| | Gant vs. Don | t = 1.088 | 20.49 | 0.8497 | mean difference = 0.08692 | -0.1446 to 0.3184 |
| | Gant vs. Bap | t = 7.367 | 18.34 | <0.0001 | mean difference = -0.3877 | -0.5418 to -0.2335 |
| | Gant vs. LaG16 | t = 5.138 | 18.93 | 0.0003 | mean difference = 0.2701 | 0.1170 to 0.4231 |
| | Don vs. Bap | t = 6.533 | 16.15 | <0.0001 | mean difference = -0.4746 | -0.6903 to -0.2589 |
| | Don vs. LaG16 | t = 2.522 | 16.25 | 0.1194 | mean difference = 0.1831 | -0.03246 to 0.3987 |
| | Bap vs. LaG16 | t = 16.16 | 19.78 | <0.0001 | mean difference = 0.6577 | 0.5398 to 0.7756 |

**Supplementary Table 5:** Comparative statistics for Figure 3. Grayed-out rows represent ANOVAs, with subsequent rows denoting multiple comparisons. ‡ represents data that were log transformed before statistical analysis.

| | Test/comparison | Statistic | df | P value | Effect size | 95% CI |
|---|---|---|---|---|---|---|
| Figure 3a | Welch's *t*-test ‡ | t = 56.95 | 23.76 | <0.0001 | mean difference = -2.120;<br>$\eta$ sqaured = 0.9927 | -2.197 to -2.043 |
| Figure 3b | Welch's *t*-test | t = 10.19 | 14.63 | <0.0001 | mean difference = 3.172;<br>$\eta$ sqaured = 0.8766 | 2.507 to 3.837 |
| Figure 3e | Welch's *t*-test | t = 23.05 | 4.013 | <0.0001 | mean difference = -32.17;<br>$\eta$ sqaured = 0.9925 | -36.04 to -28.30 |
| Figure 3g | *Ordinary one-way ANOVA with Tukey's multiple comparisons ‡* | *F = 389.5* | *3, 8* | <0.0001 | *$\eta$ squared = 0.9932* | |
| | BDNF-Ctrl vs. BDNF-Aβ | q = 40.15 | 8 | <0.0001 | mean difference = -1.133 | -1.260 to -1.005 |
| | BDNF-Ctrl vs. mCherry-Ctrl | q = 0.4317 | 8 | 0.9894 | mean difference = 0.01218 | -0.1156 to 0.1399 |
| | BDNF-Ctrl vs. mCherry-Aβ | q = 2.769 | 8 | 0.2786 | mean difference = -0.07812 | -0.2059 to 0.04963 |
| | BDNF-Aβ vs. mCherry-Ctrl | q = 40.58 | 8 | <0.0001 | mean difference = 1.145 | 1.017 to 1.272 |
| | BDNF-Aβ vs. mCherry-Aβ | q = 37.38 | 8 | <0.0001 | mean difference = 1.054 | 0.9267 to 1.182 |
| | mCherry-Ctrl vs. mCherry-Aβ | q = 3.201 | 8 | 0.1861 | mean difference = -0.0903 | -0.2180 to 0.03745 |

**Supplementary Table 6:** Comparative statistics for Figure 4. Grayed-out rows represent ANOVAs, with subsequent rows denoting multiple comparisons. ‡ represents data that were log transformed before statistical analysis.

| | Test/comparison | Statistic | df | P value | Effect size | 95% CI |
|---|---|---|---|---|---|---|
| Figure 4b | *Interaction (Cytokine x Aβ) ‡* | *F = 13.03* | *1, 15* | *0.0026* | *0.5656% of total variation* | |
| | *Cytokine ‡* | *F = 15.33* | *1, 15* | *0.0014* | *0.6656% of total variation* | |
| | *Aβ ‡* | *F = 2246* | *1, 15* | *<0.0001* | *97.47% of total variation* | |
| | +Aβ:+Cyt vs. -Aβ:+Cyt (Tukey) | q = 52.57 | 15 | <0.0001 | mean difference = 1.05 | 0.9681 to 1.131 |
| | +Aβ:+Cyt vs. +Aβ:-Cyt (Tukey) | q = 0.2975 | 15 | 0.9966 | mean difference = -0.0063 | -0.09261 to 0.08001 |
| | +Aβ:+Cyt vs. -Aβ:-Cyt (Tukey) | q = 44.81 | 15 | <0.0001 | mean difference = 0.8946 | 0.8133 to 0.9760 |
| | -Aβ:+Cyt vs. +Aβ:-Cyt (Tukey) | q = 49.86 | 15 | <0.0001 | mean difference = -1.056 | -1.142 to -0.9695 |
| | -Aβ:+Cyt vs. -Aβ:-Cyt (Tukey) | q = 7.757 | 15 | 0.0003 | mean difference = -0.1549 | -0.2363 to -0.07350 |
| | +Aβ:-Cyt vs. -Aβ:-Cyt (Tukey) | q = 42.54 | 15 | <0.0001 | mean difference = 0.9009 | 0.8146 to 0.9872 |
| Figure 4c | Interaction (Cytokine x Aβ) ‡ | *F = 17.23* | *1, 16* | *0.0008* | *0.2797% of total variation* | |
| | Cytokine ‡ | *F = 6.144* | *1, 16* | *0.0247* | *0.09974% of total variation* | |
| | Aβ ‡ | *F = 6121* | *1, 16* | *<0.0001* | *99.36% of total variation* | |
| | +Aβ:+Cyt vs. -Aβ:+Cyt (Tukey) | q = 82.39 | 16 | <0.0001 | mean difference = 2.559 | 2.433 to 2.684 |
| | +Aβ:+Cyt vs. +Aβ:-Cyt (Tukey) | q = 1.672 | 16 | 0.6461 | mean difference = 0.05193 | -0.07372 to 0.1776 |
| | +Aβ:+Cyt vs. -Aβ:-Cyt (Tukey) | q = 75.76 | 16 | <0.0001 | mean difference = 2.353 | 2.227 to 2.478 |
| | -Aβ:+Cyt vs. +Aβ:-Cyt (Tukey) | q = 80.71 | 16 | <0.0001 | mean difference = -2.507 | -2.632 to -2.381 |
| | -Aβ:+Cyt vs. -Aβ:-Cyt (Tukey) | q = 6.63 | 16 | 0.0013 | mean difference = -0.2059 | -0.3315 to -0.08023 |
| | +Aβ:-Cyt vs. -Aβ:-Cyt (Tukey) | q = 74.08 | 16 | <0.0001 | mean difference = 2.301 | 2.175 to 2.426 |
| Figure 4d *IL6* | *Interaction (Cyt × Aβ)* | *F = 33.80* | *1, 16* | *<0.0001* | *7.919% of total variation* | |
| | *Cytokine* | *F = 369.4* | *1, 16* | *<0.0001* | *86.53% of total variation* | |
| | *Aβ* | *F = 7.681* | *1, 16* | *0.0136* | *1.799% of total variation* | |
| | +Aβ:+Cyt vs. -Aβ:+Cyt (Tukey) | q = 8.585 | 16.000 | <0.0001 | mean difference = 1.710 | 0.9041 to 2.516 |
| | +Aβ:+Cyt vs. +Aβ:-Cyt (Tukey) | q = 13.41 | 16.000 | <0.0001 | mean difference = -2.670 | -3.476 to -1.864 |
| | +Aβ:+Cyt vs. -Aβ:-Cyt (Tukey) | q = 16.45 | 16.000 | <0.0001 | mean difference = -3.276 | -4.082 to -2.470 |

| | | | | | | |
|---|---|---|---|---|---|---|
| | -Aβ:+Cyt vs. +Aβ:-Cyt (Tukey) | q = 21.99 | 16.000 | <0.0001 | mean difference = -4.380 | -5.186 to -3.574 |
| | -Aβ:+Cyt vs. -Aβ:-Cyt (Tukey) | q = 25.03 | 16.000 | <0.0001 | mean difference = -4.986 | -5.792 to -4.180 |
| | +Aβ:-Cyt vs. -Aβ:-Cyt (Tukey) | q = 3.042 | 16.000 | 0.1794 | mean difference = -0.6060 | -1.412 to 0.1999 |
| Figure 4d *C3* | *Interaction (Cyt × Aβ)* | *F = 21.33* | *1, 16* | *0.0003* | *2.903% of total variation* | |
| | *Cytokine* | *F = 679.2* | *1, 16* | *<0.0001* | *92.47% of total variation* | |
| | *Aβ* | *F = 17.97* | *1, 16* | *0.0006* | *2.446% of total variation* | |
| | +Aβ:+Cyt vs. -Aβ:+Cyt (Tukey) | q = 8.857 | 16.000 | <0.0001 | mean difference = 1.776 | 0.9647 to 2.587 |
| | +Aβ:+Cyt vs. +Aβ:-Cyt (Tukey) | q = 21.44 | 16.000 | <0.0001 | mean difference = -4.300 | -5.111 to -3.489 |
| | +Aβ:+Cyt vs. -Aβ:-Cyt (Tukey) | q = 21.82 | 16.000 | <0.0001 | mean difference = -4.376 | -5.187 to -3.565 |
| | -Aβ:+Cyt vs. +Aβ:-Cyt (Tukey) | q = 30.30 | 16.000 | <0.0001 | mean difference = -6.076 | -6.887 to -5.265 |
| | -Aβ:+Cyt vs. -Aβ:-Cyt (Tukey) | q = 30.68 | 16.000 | <0.0001 | mean difference = -6.152 | -6.963 to -5.341 |
| | +Aβ:-Cyt vs. -Aβ:-Cyt (Tukey) | q = 0.3790 | 16.000 | 0.9930 | mean difference = -0.07600 | -0.8873 to 0.7353 |
| Figure 4d *CSF* | *Interaction (Cyt × Aβ)* | *F = 50.00* | *1, 15* | *<0.0001* | *9.049% of total variation* | |
| | *Cytokine* | *F = 495.3* | *1, 15* | *<0.0001* | *89.63% of total variation* | |
| | *Aβ* | *F = 21.34* | *1, 15* | *0.0003* | *3.862% of total variation* | |
| | +Aβ:+Cyt vs. -Aβ:+Cyt (Tukey) | q = 11.36 | 15.000 | <0.0001 | mean difference = 2.661 | 1.706 to 3.616 |
| | +Aβ:+Cyt vs. +Aβ:-Cyt (Tukey) | q = 15.65 | 15.000 | <0.0001 | mean difference = -3.456 | -4.356 to -2.556 |
| | +Aβ:+Cyt vs. -Aβ:-Cyt (Tukey) | q = 18.18 | 15.000 | <0.0001 | mean difference = -4.014 | -4.914 to -3.114 |
| | -Aβ:+Cyt vs. +Aβ:-Cyt (Tukey) | q = 26.12 | 15.000 | <0.0001 | mean difference = -6.117 | -7.072 to -5.162 |
| | -Aβ:+Cyt vs. -Aβ:-Cyt (Tukey) | q = 28.50 | 15.000 | <0.0001 | mean difference = -6.675 | -7.630 to -5.720 |
| | +Aβ:-Cyt vs. -Aβ:-Cyt (Tukey) | q = 2.527 | 15.000 | 0.3172 | mean difference = -0.5580 | -1.458 to 0.3420 |
| Figure 4e *IL6* | *Interaction (Cyt × Aβ)* | *F = 1.925* | *1, 16* | *0.1843* | *0.2138% of total variation* | |
| | *Cytokine* | *F = 878.6* | *1, 16* | *<0.0001* | *97.59% of total variation* | |
| | *Aβ* | *F = 3.773* | *1, 16* | *0.0699* | *0.4191% of total variation* | |
| | +Aβ:+Cyt vs. -Aβ:+Cyt (Tukey) | q = 0.5550 | 16.000 | 0.9788 | mean difference = -0.08800 | -0.7296 to 0.5536 |

| | | | | | | |
|---|---|---|---|---|---|---|
| | +Aβ:+Cyt vs. +Aβ:-Cyt (Tukey) | q = 28.25 | 16.000 | <0.0001 | mean difference = -4.480 | -5.122 to -3.838 |
| | +Aβ:+Cyt vs. -Aβ:-Cyt (Tukey) | q = 31.58 | 16.000 | <0.0001 | mean difference = -5.008 | -5.650 to -4.366 |
| | -Aβ:+Cyt vs. +Aβ:-Cyt (Tukey) | q = 27.70 | 16.000 | <0.0001 | mean difference = -4.392 | -5.034 to -3.750 |
| | -Aβ:+Cyt vs. -Aβ:-Cyt (Tukey) | q = 31.03 | 16.000 | <0.0001 | mean difference = -4.920 | -5.562 to -4.278 |
| | +Aβ:-Cyt vs. -Aβ:-Cyt (Tukey) | q = 3.330 | 16.000 | 0.1270 | mean difference = -0.5280 | -1.170 to 0.1136 |
| Figure 4e *C3* | *Interaction (Cyt × Aβ)* | F = 1.001 | 1, 15 | 0.3329 | 0.02692% of total variation | |
| | *Cytokine* | F = 3679 | 1, 15 | <0.0001 | 98.96% of total variation | |
| | *Aβ* | F = 0.4725 | 1, 15 | 0.5023 | 0.01271% of total variation | |
| | +Aβ:+Cyt vs. -Aβ:+Cyt (Tukey) | q = 1.740 | 15.000 | 0.6182 | mean difference = -0.1860 | -0.6217 to 0.2497 |
| | +Aβ:+Cyt vs. +Aβ:-Cyt (Tukey) | q = 59.92 | 15.000 | <0.0001 | mean difference = -6.795 | -7.257 to -6.332 |
| | +Aβ:+Cyt vs. -Aβ:-Cyt (Tukey) | q = 63.23 | 15.000 | <0.0001 | mean difference = -6.760 | -7.196 to -6.324 |
| | -Aβ:+Cyt vs. +Aβ:-Cyt (Tukey) | q = 58.28 | 15.000 | <0.0001 | mean difference = -6.609 | -7.071 to -6.146 |
| | -Aβ:+Cyt vs. -Aβ:-Cyt (Tukey) | q = 61.49 | 15.000 | <0.0001 | mean difference = -6.574 | -7.010 to -6.138 |
| | +Aβ:-Cyt vs. -Aβ:-Cyt (Tukey) | q = 0.3043 | 15.000 | 0.9963 | mean difference = 0.03450 | -0.4277 to 0.4967 |
| Figure 4e *CSF* | *Interaction (Cyt × Aβ)* | F = 1.665 | 1, 16 | 0.2152 | 0.2491% of total variation | |
| | *Cytokine* | F = 650.8 | 1, 16 | <0.0001 | 97.35% of total variation | |
| | *Aβ* | F = 0.07549 | 1, 16 | 0.7870 | 0.01129% of total variation | |
| | +Aβ:+Cyt vs. -Aβ:+Cyt (Tukey) | q = 1.016 | 16.000 | 0.8884 | mean difference = -0.2440 | -1.216 to 0.7280 |
| | +Aβ:+Cyt vs. +Aβ:-Cyt (Tukey) | q = 26.80 | 16.000 | <0.0001 | mean difference = -6.438 | -7.410 to -5.466 |
| | +Aβ:+Cyt vs. -Aβ:-Cyt (Tukey) | q = 25.24 | 16.000 | <0.0001 | mean difference = 6.062 | -7.034 to -5.090 |
| | -Aβ:+Cyt vs. +Aβ:-Cyt (Tukey) | q = 25.78 | 16.000 | <0.0001 | mean difference = -6.194 | -7.166 to -5.222 |
| | -Aβ:+Cyt vs. -Aβ:-Cyt (Tukey) | q = 24.22 | 16.000 | <0.0001 | mean difference = -5.818 | -6.790 to -4.846 |
| | +Aβ:-Cyt vs. -Aβ:-Cyt (Tukey) | q = 1.565 | 16.000 | 0.6906 | mean difference = 0.3760 | -0.5960 to 1.348 |

**Supplementary Table 7:** Comparative statistics for Figure 5. Grayed-out rows represent ANOVAs, with subsequent rows denoting multiple comparisons. ‡ represents data that were log transformed before statistical analysis.

| | Test/comparison | Statistic | df | P value | Effect size | 95% CI |
|---|---|---|---|---|---|---|
| Figure 5b | *Welch ANOVA with Dunnett's T3 multiple comparisons ‡* | *W = 24.83* | *5.000, 13.98* | *<0.0001* | | |
| | WT JMB525 vs. WT JCL8 | t = 2.143 | 15.63 | 0.4452 | mean difference = 0.08881 | -0.05114 to 0.2288 |
| | WT JMB525 vs. WT No cell | t = 7.623 | 4.47 | 0.0125 | mean difference = 0.3448 | 0.1112 to 0.5785 |
| | WT JMB525 vs. Hemi JMB525 | t = 6.447 | 11.45 | 0.0006 | mean difference = -0.6641 | -1.035 to -0.2931 |
| | WT JMB525 vs. Hemi JCL8 | t = 3.949 | 16.71 | 0.0143 | mean difference = 0.1922 | 0.02920 to 0.3552 |
| | WT JMB525 vs. Hemi No cell | t = 6.589 | 7.905 | 0.0021 | mean difference = 0.3104 | 0.1271 to 0.4936 |
| | WT JCL8 vs. WT No cell | t = 5.326 | 5.374 | 0.0278 | mean difference = 0.256 | 0.03440 to 0.4777 |
| | WT JCL8 vs. Hemi JMB525 | t = 7.22 | 11.96 | 0.0001 | mean difference = -0.7529 | -1.122 to -0.3837 |
| | WT JCL8 vs. Hemi JCL8 | t = 2.015 | 17.67 | 0.5246 | mean difference = 0.1034 | -0.06719 to 0.2740 |
| | WT JCL8 vs. Hemi No cell | t = 4.446 | 9.076 | 0.0192 | mean difference = 0.2216 | 0.03366 to 0.4095 |
| | WT No cell vs. Hemi JMB525 | t = 9.532 | 11.8 | <0.0001 | mean difference = -1.009 | -1.384 to -0.6342 |
| | WT No cell vs. Hemi JCL8 | t = 2.805 | 7.693 | 0.2136 | mean difference = -0.1526 | -0.3644 to 0.05911 |
| | WT No cell vs. Hemi No cell | t = 0.65 | 5.539 | 0.9996 | mean difference = -0.03447 | -0.2612 to 0.1922 |
| | Hemi JMB525 vs. Hemi JCL8 | t = 7.976 | 13.23 | <0.0001 | mean difference = 0.8563 | 0.4817 to 1.231 |
| | Hemi JMB525 vs. Hemi No cell | t = 9.136 | 12.53 | <0.0001 | mean difference = 0.9745 | 0.6023 to 1.347 |
| | Hemi JCL8 vs. Hemi No cell | t = 2.111 | 11.89 | 0.4776 | mean difference =0.1182 | -0.08005 to 0.3164 |

**Supplementary Table 8:** Comparative statistics for Figure 6. Grayed-out rows represent ANOVAs, with subsequent rows denoting multiple comparisons.

| | Test/comparison | Statistic | df | P value | Effect size |
|---|---|---|---|---|---|
| Figure 6b | *Kruskal-Wallis with Dunn's multiple comparisons test* | *H = 7.821* | *3* | *0.0205* | |
| | 5xFAD Bap-Notch vs. WT Bap-Notch | Z = 2.717 | 3 | 0.0395 | mean rank difference = 8 |
| | 5xFAD Bap-Notch vs. 5xFAD Uninjected | Z = 1.925 | 3 | 0.3255 | mean rank difference = 5.667 |
| | 5xFAD Bap-Notch vs. WT Uninjected | Z = 1.472 | 3 | 0.8462 | mean rank difference = 4.333 |
| | WT Bap-Notch vs. 5xFAD Uninjected | Z = 0.7926 | 3 | >0.9999 | mean rank difference = -2.333 |
| | WT Bap-Notch vs. WT Uninjected | Z = 1.246 | 3 | >0.9999 | mean rank difference = -3.667 |
| | 5xFAD Uninjected vs. WT Uninjected | Z = 0.4529 | 3 | >0.9999 | mean rank difference = -1.333 |
| Figure 6c | Mann-Whitney test | U = 4 | | >0.9999 | difference between medians = -62.00 |

**Supplementary Table 9:** Comparative statistics for Supplementary Figure 2. Grayed-out rows represent ANOVAs, with subsequent rows denoting multiple comparisons. ‡ represents data that were log transformed before statistical analysis.

| | **Test/comparison** | **Statistic** | **df** | **P value** | **Effect size** | **95% CI** |
|---|---|---|---|---|---|---|
| Supp Figure 2b | *Ordinary one-way ANOVA with Tukey's multiple comparisons* | *F = 474.4* | *2, 6* | *<0.0001* | *η squared = 0.9937* | |
| | Aβ vs. Control | q = 0.5798 | 6 | 0.9128 | mean difference = -6756 | -57319 to 43807 |
| | Aβ vs. Myc | q = 38.03 | 6 | <0.0001 | mean difference = -443090 | -493653 to -392527 |
| | Control vs. Myc | q = 37.45 | 6 | <0.0001 | mean difference = -436334 | -486897 to -385771 |
| Supp Figure 2c | *Ordinary one-way ANOVA with Tukey's multiple comparisons ‡* | *F = 4173* | *2, 6* | *<0.0001* | *η squared = 0.9993* | |
| | Aβ vs. Control | q = 0.1211 | 6 | 0.996 | mean difference = 0.002196 | -0.07652 to 0.08091 |
| | Aβ vs. Myc | q = 111.8 | 6 | <0.0001 | mean difference = -2.029 | -2.107 to -1.950 |
| | Control vs. Myc | q = 112 | 6 | <0.0001 | mean difference = -2.031 | -2.110 to -1.952 |

**Supplementary Table 10:** Comparative statistics for Supplementary Figure 4. Grayed-out rows represent ANOVAs, with subsequent rows denoting multiple comparisons.

| | Test/comparison | Statistic | df | P value | Effect size | 95% CI |
|---|---|---|---|---|---|---|
| Supp Figure 4a | *Interaction (Cytokine × Aβ)* | *F = 4.192* | *1, 16* | *0.0574* | *11.66% of total variation* | |
| | *Cytokine* | *F = 15.58* | *1, 16* | *0.0012* | *43.33% of total variation* | |
| | *Aβ* | *F = 0.1850* | *1, 16* | *0.6729* | *0.5144% of total variation* | |
| | +Aβ:+Cyt vs. -Aβ:+Cyt (Tukey) | q = 2.478 | 16 | 0.331 | mean difference = 26.7 | -16.90 to 70.31 |
| | +Aβ:+Cyt vs. +Aβ:-Cyt (Tukey) | q = 1.9 | 16 | 0.5503 | mean difference = -20.48 | -64.08 to 23.13 |
| | +Aβ:+Cyt vs. -Aβ:-Cyt (Tukey) | q = 3.517 | 16 | 0.1003 | mean difference = -37.91 | -81.52 to 5.698 |
| | -Aβ:+Cyt vs. +Aβ:-Cyt (Tukey) | q = 4.378 | 16 | 0.0317 | mean difference = -47.18 | -90.79 to -3.573 |
| | -Aβ:+Cyt vs. -Aβ:-Cyt (Tukey) | q = 5.995 | 16 | 0.0031 | mean difference = -64.61 | -108.2 to -21.00 |
| | +Aβ:-Cyt vs. -Aβ:-Cyt (Tukey) | q = 1.617 | 16 | 0.669 | mean difference = -17.43 | -61.04 to 26.18 |
| Supp Figure 4c | *Interaction (Cytokine × Aβ)* | *F = 5.802* | *1, 16* | *0.0284* | *6.878% of total variation* | |
| | *Cytokine* | *F = 4.906* | *1, 16* | *0.0416* | *5.816% of total variation* | |
| | *Aβ* | *F = 57.65* | *1, 16* | *<0.0001* | *68.34% of total variation* | |
| | +Aβ:+Cyt vs. -Aβ:+Cyt (Tukey) | q = 10 | 16 | <0.0001 | mean difference = 2570 | 1530 to 3610 |
| | +Aβ:+Cyt vs. +Aβ:-Cyt (Tukey) | q = 4.624 | 16 | 0.0225 | mean difference = 1188 | 148.5 to 2228 |
| | +Aβ:+Cyt vs. -Aβ:-Cyt (Tukey) | q = 9.808 | 16 | <0.0001 | mean difference = 2521 | 1481 to 3560 |
| | -Aβ:+Cyt vs. +Aβ:-Cyt (Tukey) | q = 5.378 | 16 | 0.0077 | mean difference = -1382 | -2422 to -342.2 |
| | -Aβ:+Cyt vs. -Aβ:-Cyt (Tukey) | q = 0.1938 | 16 | 0.999 | mean difference = -49.8 | -1090 to 990.0 |
| | +Aβ:-Cyt vs. -Aβ:-Cyt (Tukey) | q = 5.184 | 16 | 0.0101 | mean difference = 1332 | 292.4 to 2372 |
| | *Interaction (Cytokine × Aβ)* | *F = 0.5720* | *1, 16* | *0.4604* | *0.01276% of total variation* | |

| | | | | | | |
|---|---|---|---|---|---|---|
| Supp Figure 4d | *Cytokine* | *F = 16.32* | *1, 16* | *0.0009* | *0.3641% of total variation* | |
| | *Aβ* | *F = 4451* | *1, 16* | *<0.0001* | *99.27% of total variation* | |
| | +Aβ:+Cyt vs. -Aβ:+Cyt (Tukey) | q = 65.96 | 16 | <0.0001 | mean difference = 22739230 | 21344320 to 24134139 |
| | +Aβ:+Cyt vs. +Aβ:-Cyt (Tukey) | q = 4.797 | 16 | 0.0176 | mean difference = -1653637 | -3048546 to -258727 |
| | +Aβ:+Cyt vs. -Aβ:-Cyt (Tukey) | q = 62.67 | 16 | <0.0001 | mean difference = 21607087 | 20212177 to 23001997 |
| | -Aβ:+Cyt vs. +Aβ:-Cyt (Tukey) | q = 70.75 | 16 | <0.0001 | mean difference = -24392866 | -25787776 to -22997957 |
| | -Aβ:+Cyt vs. -Aβ:-Cyt (Tukey) | q = 3.284 | 16 | 0.1344 | mean difference = -1132143 | -2527052 to 262767 |
| | +Aβ:-Cyt vs. -Aβ:-Cyt (Tukey) | q = 67.47 | 16 | <0.0001 | mean difference = 23260724 | 21865814 to 24655633 |

**Supplementary Table 11:** Comparative statistics for Supplementary Figure 6.

| | **Test/comparison** | **Statistic** | **df** | **P value** | **Effect size** | **95% CI** |
|---|---|---|---|---|---|---|
| Supp Figure 6a | Welch's *t*-test | t = 1.84 | 2.294 | 0.19 | mean difference = -50.46; η squared = 0.5971 | -154.8 to 53.92 |
| Supp Figure 6b | Welch's *t*-test | t = 2.909 | 2.52 | 0.0766 | mean difference = 13.96; η squared = 0.7704 | -3.097 to 31.01 |
| Supp Figure 6c | Welch's *t*-test | t = 0.1762 | 2.62 | 0.8728 | mean difference = 4.044; η squared = 0.01171 | -75.36 to 83.45 |
| Supp Figure 6d | Welch's *t*-test | t=0.1933 | 3.599 | 0.8572 | mean difference = 0.01955; η squared = 0.01027 | -0.2741 to 0.3132 |
| Supp Figure 6e | Welch's *t*-test | t = 2.302 | 2.113 | 0.1411 | mean difference = 47.95; η squared = 0.7150 | -37.22 to 133.1 |
| Supp Figure 6f | Welch's *t*-test | t=1.190 | 2.562 | 0.3325 | mean difference = -0.1736; η squared = 0.3561 | -0.6858 to 0.3387 |

**Supplementary Table 12:** Comparative statistics for Supplementary Figure 7. Grayed-out rows represent ANOVAs, with subsequent rows denoting multiple comparisons.

| | **Test/comparison** | **Statistic** | **df** | **P value** | **Effect size** | **95% CI** |
|---|---|---|---|---|---|---|
| Supp Figure 7a | *Welch ANOVA with Dunnett's T3 multiple comparisons* | W = 1520 | 2.000, 4.493 | 0.0003 | | |
| | Aβ42 vs Aβ40 | t = 1.9 | 3.382 | 0.3294 | mean difference = 0.4564 | -0.6079 to 1.521 |
| | Aβ42 vs Control | t = 57.9 | 3.787 | <0.0001 | mean difference = 3.631 | 3.397 to 3.866 |
| | Aβ40 vs Control | t = 13.57 | 3.051 | 0.0021 | mean difference = 3.175 | 2.139 to 4.211 |
| Supp Figure 7b | *Welch ANOVA with Dunnett's T3 multiple comparisons* | W = 45.07 | 2.000, 5.350 | 0.0004 | | |
| | Aβ42 vs Aβ40 (Tukey) | t = 0.8454 | 5.848 | 0.7853 | mean difference = 1.6 | -4.442 to 7.641 |
| | Aβ42 vs Control (Tukey) | t = 8.736 | 4.644 | 0.0009 | mean difference = 12.2 | 7.452 to 16.94 |
| | Aβ40 vs Control (Tukey) | t = 6.666 | 4.236 | 0.0067 | mean difference = 10.6 | 4.645 to 16.55 |
| Supp Figure 7c | *Welch ANOVA with Dunnett's T3 multiple comparisons* | W = 19.15 | 2.000, 4.016 | 0.0088 | | |
| | Aβ42 vs Aβ40 (Tukey) | t = 1.242 | 4.734 | 0.5628 | mean difference = -7.078 | -26.46 to 12.30 |
| | Aβ42 vs Control (Tukey) | t = 5.112 | 3.018 | 0.0341 | mean difference = 14.34 | 1.914 to 26.77 |
| | Aβ40 vs Control (Tukey) | t = 4.312 | 3.006 | 0.0537 | mean difference = 21.42 | -0.5847 to 43.42 |
